# Molecular design principles for Photosystem I-based biohybrid solar fuel catalysts

**DOI:** 10.64898/2026.03.23.713776

**Authors:** Maximino D. Emerson, Siva Naga Sai Damaraju, Audrey H. Short, Zachary B. Alvord, Zsolt A. Palmer, Himanshu S. Mehra, Christian M. Brininger, Josh V. Vermaas, Lisa M. Utschig, Christopher J. Gisriel

## Abstract

Direct solar-to-chemical conversion offers a compelling route to clean, dispatchable energy. Photosystem I (PSI), an evolutionarily optimized light-driven oxidoreductase central to oxygenic photosynthesis, can be repurposed for direct solar-fuel production by efficiently coupling its photochemistry to catalysts, thereby storing sunlight as chemical energy in the H-H bond of H2. One promising architecture integrates PSI with Pt nanoparticle (PtNP) catalysts to create photocatalytic PSI-PtNP biohybrids. Advancing these systems requires molecular-level insight into protein-nanoparticle interactions and the bio-nano electron transfer pathways that govern activity; however, progress has been constrained by limited structural data to guide rational design. Here, we present two molecular structures of active PSI-PtNP assemblies that (a) compare thermophilic and mesophilic PSI scaffolds and (b) probe how removal of the terminal [4Fe-4S] clusters and stromal subunits in PSI reshapes protein-nanoparticle interfaces and photocatalysis. Structural analyses and molecular dynamics simulations define the interface topology, electrostatics, and cofactor-to-nanoparticle distances, revealing key molecular features that control biohybrid formation and electron transfer efficiency. These data establish mechanistic links between scaffold composition, bio-nano interface geometry, and catalytic performance, yielding design principles for optimizing PSI-PtNP architectures. The resulting structure-function insights provide a blueprint for engineering PSI-based solar-fuels systems and, more broadly, inform the design of protein-nanomaterial interfaces for light-driven catalysis.

## Introduction

Photosynthetic organisms capture sunlight and convert it into chemical energy that fuels the synthesis of biomolecules, thereby storing much of the energy that enters Earth’s biosphere. In oxygenic photosynthesis, this initial conversion is performed by two large transmembrane pigment-protein complexes, Photosystem II (PSII) and Photosystem I (PSI). These proteins effectively capture and convert sunlight into usable electrochemical potential via a series of light-induced rapid, sequential electron transfer reactions. This light-driven redox chemistry powers the core reactions of photosynthesis and generates reducing equivalents and ATP that ultimately drive the reduction of CO_2_ to organic carbon. Leveraging this activity to instead produce valuable fuels has been pursued as a method of meeting increasing global energy needs without depending on limited geological resources or the environmental impact associated with mining and burning fossil fuels. One promising photons-to-fuels strategy uses PSI to harvest solar energy and platinum nanoparticles (PtNPs) to generate H_2_, converting solar energy to a stable, storable form.

Cyanobacterial PSI is predominantly found as an ∼1 MDa trimer of ∼330 kDa monomers. Each monomer consists of 10-12 protein subunits and coordinates ∼95 light-harvesting chlorophyll molecules that absorb and transfer energy. At the core of each monomer are two pseudo-C2 symmetric electron transfer branches made up of embedded donor and acceptor cofactors that span the membrane (*SI Appendix*, **Fig. S1**). The sequence of reactions begins near the lumenal side of the complex with a special chlorophyll *a/*chlorophyll *a*′ pair called P_700_, followed by four monomeric chlorophyll *a* molecules (A_−1A_, A_−1B_, A_0A_, and A_0B_) (1), then two quinones (A_1A_ and A_1B_) (2), and concludes on the stromal side with three [4Fe-4S] clusters (F_X_, F_A_, and F_B_) (3). Following photoexcitation of PSI, P_700_ donates an electron into one of the two branches where it is transferred sequentially through the A_0_, A_1_, F_X_, F_A_ and F_B_ cofactors. This spatially separates the electron and hole to limit charge recombination and transfers the light-generated reducing equivalents from the inaccessible core of PSI to surface sites accessible to redox protein acceptors ferredoxin (Fd) or flavodoxin (Flv). For each step further down the chain of redox cofactors, the lifetime of the charge-separated state increases. The final charge-separated state, P_700_^+^F_B_ ^−^, is stable for ∼65 ms (4), which is sufficient time for the electron to be transferred to a native acceptor or an abiotic catalyst in PSI biohybrids. Additionally, the broad absorption profile of PSI, near-unity quantum efficiency, and strongly negative redox potential of the terminal acceptor F_B_ (−580 mV vs. NHE, pH independent) (5) make PSI an ideal platform for developing systems that use light to generate H_2_ from water (E_m_, pH 6.3 = −370 mV) (6).

Considerable effort has been devoted to constructing PSI-biohybrid catalysts that efficiently redirect PSI photochemistry toward H_2_ evolution. In seminal work, Greenbaum and co-workers photochemically deposited metallic Pt onto chloroplast thylakoid membranes, achieving light-driven concurrent H_2_ and O_2_ evolution (7). Building on this foundation, photoprecipitation of metallic Pt onto isolated PSI has continued to be investigated (8–14). In subsequent systems, photogenerated electrons from PSI have been coupled to hydrogenases (15–19), platinum catalysts (20–27), and molecular H_2_-evolution catalysts (28–30). A robust approach to forming PSI biohybrids uses electrostatically driven self-assembly: negatively charged PtNPs bind to the stromal face of PSI, mimicking native acceptor-protein interactions (20). This strategy yields discrete complexes rather than ill-defined mixtures, facilitating isolation, rigorous electron paramagnetic resonance spectroscopic characterization, and cryogenic electron microscopy (cryo-EM) structure determination. Moreover, the same targeted self-assembly strategy translates to PtNP incorporation at intrinsic PSI sites within thylakoid membranes, enabling complete solar water-splitting systems (31, 32).

The PSI-PtNP biohybrids were first generated using PSI from the cyanobacteria *Synechococcus lividus* PCC 6717 (hereafter *S. lividus*) and *Synechococcus leopoliensis*; however, a lack of structural data hindered the ability to identify molecular features underlying PtNP binding and photocatalytic activity. To fill this gap, a recent PSI-PtNP biohybrid structure was determined using cryo-EM which contained PSI isolated from *S. lividus* (33). In each monomer of the trimeric PSI holocomplex, the structure showed one PtNP binding site near the terminal side of the electron transfer chain in a similar location to where native acceptors Fd and Flv bind, and an additional distal site too far from the electron transfer chain to be involved in H_2_ production. While this structure revealed the locations of PtNP binding to trimeric PSI in that species, questions persist, such as whether PtNPs bind similarly to PSI from other organisms and exactly what interactions govern PtNP binding.

In PSI, forward electron transfer to the terminal [4Fe-4S] clusters, F_A_ and F_B_, overwhelmingly outcompetes charge recombination with P_700_^+^, resulting in near-stoichiometric delivery to these sites. Yet, the most reducing (highest-energy) electrons reside earlier along the electron transfer chain, where – if intercepted before recombination – they drive more demanding chemistry. To access these higher-energy electrons at F_X_ (≈ −700 mV vs NHE) (2), we engineered a PSI core lacking stromal subunits PsaC, PsaD, and PsaE, the former of which coordinates F_A_ and F_B_, to intercept electrons from the intermediate charge-separated state P_700_^+^F_X_ ^−^ (lifetime ≈ 1 ms) (34). We show that targeted PtNP coupling to the PSI core biohybrid facilitates electron capture from the intermediate charge-separated state and supports H_2_ evolution. We also determine the structure of a PSI-PtNP biohybrid from *Thermosynechococcus vestitus*, a model thermophilic cyanobacterium. Structural and molecular dynamics (MD) comparisons of the PSI core biohybrid to the native trimer biohybrid – whose surface topology, charge, and distance to the nearest cofactors differ – helps to clarify poorly understood mechanisms of electron exchange at bio-nano interfaces and reveal how interface design biases electron transfer efficiency and charge accumulation at PtNP catalyst sites. These results reveal new PtNP-PSI interactions useful for engineering biohybrid bio-nano interfaces for efficient electron transfer and photocatalytic function.

## Results

### Preparation and characterization of PSI-PtNP biohybrids

PSI was isolated from *T. vestitus* and *S. leopoliensis* as described in **Materials and Methods**, each of which exhibited a mixed chlorophyll absorption spectrum typical of PSI isolations with Q_x_ and Q_y_ transition maxima centered at ∼595 and ∼680 nm, respectively (*SI Appendix*, **Fig. S2A**). Room temperature fluorescence was also typical of PSI isolations; *S. leopoliensis* PSI exhibits a single fluorescence maximum at ∼680 nm and *T. vestitus* PSI exhibits two maxima at 688 nm and 715 nm, the latter of which is due to low energy chlorophyll sites (35) (*SI Appendix*, **Fig. S2B**). For *T. vestitus* PSI, size exclusion chromatography (*SI Appendix*, **Fig. S3**) showed a peak maximum for *T. vestitus* PSI eluting at ∼7.0 min. A peak maximum is also observed at ∼7.0 min for *S. leopoliensis* PSI, but there is an additional shoulder centered at ∼8.0 min. We hypothesized this to indicate that PSI from both organisms is primarily in a trimeric oligomeric state but that *S. leopoliensis* PSI contains a small fraction of monomeric PSI.

PSI cores lacking the stromal subunits and terminal [4Fe-4S] clusters F_A_ and F_B_ were generated from *S. leopoliensis* PSI as previously established using treatment with 6.8 M urea (36). Although this procedure was developed in the 1980s, the oligomeric state of the sample was not reported. Size-exclusion chromatography of urea-treated PSI showed a peak maximum at ∼8.3 min, and an additional shoulder at ∼7.0 min, that accounts for ∼5% of the peak area (*SI Appendix*, **Fig. S3**). We hypothesized this to indicate that the urea-treated *S. leopoliensis* PSI are present primarily as monomeric cores except for a minor fraction that remain assembled as trimeric complexes.

PtNPs of ∼2 nm in diameter were synthesized and bound to the PSI samples as described previously (33). Photocatalytic H_2_ production was measured by gas chromatography. Under 3,000 μE illumination, the biohybrids using trimeric PSI isolated from *T. vestitus* generated H_2_ at a maximal turnover frequency of 5,360 mol H_2_ mol PSI^−1^ h^−1^ and had a turnover number of 31,500 in 480 min. The biohybrids using PSI cores generated from *S. leopoliensis* PSI produced H_2_ at a maximal turnover frequency of 1,300 mol H_2_ mol PSI^−1^ h^−1^ and had a turnover number of 2,100 in 168 minutes.

Each biohybrid sample was negatively stained and imaged by transmission electron microscopy (TEM) as described in **Materials and Methods**. Consistent with our interpretation of the size exclusion chromatography data, biohybrids using PSI isolated from *T. vestitus* showed a typical trimeric configuration (*SI Appendix*, **Fig. S4**), and biohybrids using urea-treated PSI isolated from *S. leopoliensis* appeared as monomeric complexes (*SI Appendix*, **Fig. S4**). Hereafter, we refer to the two PtNP biohybrid samples as Pt-PSI^tri^_*T*.*v*._, and Pt-PSI^cor^_*S*.*leo*._.

Because no structural data yet exists for PSI from *S. leopoliensis*, we also generated sequence alignments and calculated sequence identities with PSI subunits from *T. vestitus, S. lividus* (from which a previous PSI-PtNP biohybrid structure was determined) (33), and *Synechocystis* sp. PCC 6803 (hereafter *Synechocystis* 6803) (*SI Appendix*, **Fig. S5, Fig. S6**, and **Table S1**). Sequence identities for different subunits span from ∼90% (e.g., PsaA, PsaB, and PsaC) to ∼45% (e.g., PsaL).

### Structural determination of PSI-PtNP biohybrids

Single particle cryo-EM was performed on the Pt-PSI^tri^_*T*.*v*._ and Pt-PSI^cor^_*S*.*leo*._ samples and data were processed as described in **Material and Methods**. The Pt-PSI^tri^_*T*.*v*._ structure was determined to 3.39 Å global resolution (*SI Appendix*, **Fig. S7**). The corresponding data statistics and processing workflow can be found in *SI Appendix*, **Table S2** and **Fig. S8**. The structure of Pt-PSI^tri^_*T*.*v*._ is similar to the previously reported biohybrid structure of trimeric *S. lividus* PSI with bound PtNPs (33) (hereafter Pt-PSI^tri^_*S*.*liv*._). Each monomer of the trimer contains subunits PsaA, PsaB, PsaC, PsaD, PsaE, PsaF, PsaI, PsaJ, PsaK, PsaL, PsaM, and PsaX that coordinate 285 chlorophyll *a* molecules, 3 chlorophyll *a′* molecules, 66 β-carotenes, 6 phylloquinones, 9 [4Fe-4S] clusters, and 9 lipids. On each monomer, two high-signal, irregular ellipsoid regions were assigned as PtNP binding sites (**Fig. 1**). The site closest to the native electron transfer cofactors (∼14 Å from the [4Fe-4S] cluster F_B_) is hereafter referred to as site A while the site more distal (∼38 Å from F_B_) is referred to as site B. Site A is positioned near subunits PsaA, PsaE, PsaC, PsaF, and PsaJ, overlapping somewhat the region where native electron acceptors bind (37–40). This signal is ∼40 Å at its longest dimension by 34 Å wide by 20 Å thick. Site B is positioned at the periphery of the monomer near subunits PsaB, PsaE, PsaF, and PsaX. The signal at site B is more spherical, ∼30 Å in its longest dimension and 20 Å in both orthogonal dimensions. Like the Pt-PSI^tri^_*S*.*liv*._ structure (33), both PtNP sites show potential interactions with the detergent micelle, lack discrete signals for individual atoms, and exhibit irregular ellipsoidal shapes and ill-defined edges of the signal. Thus, although the PtNPs generally bind in sites A and B, there is heterogeneity in the exact positions, orientations, and probably extents to which the sites are occupied.

**Fig. 1.**
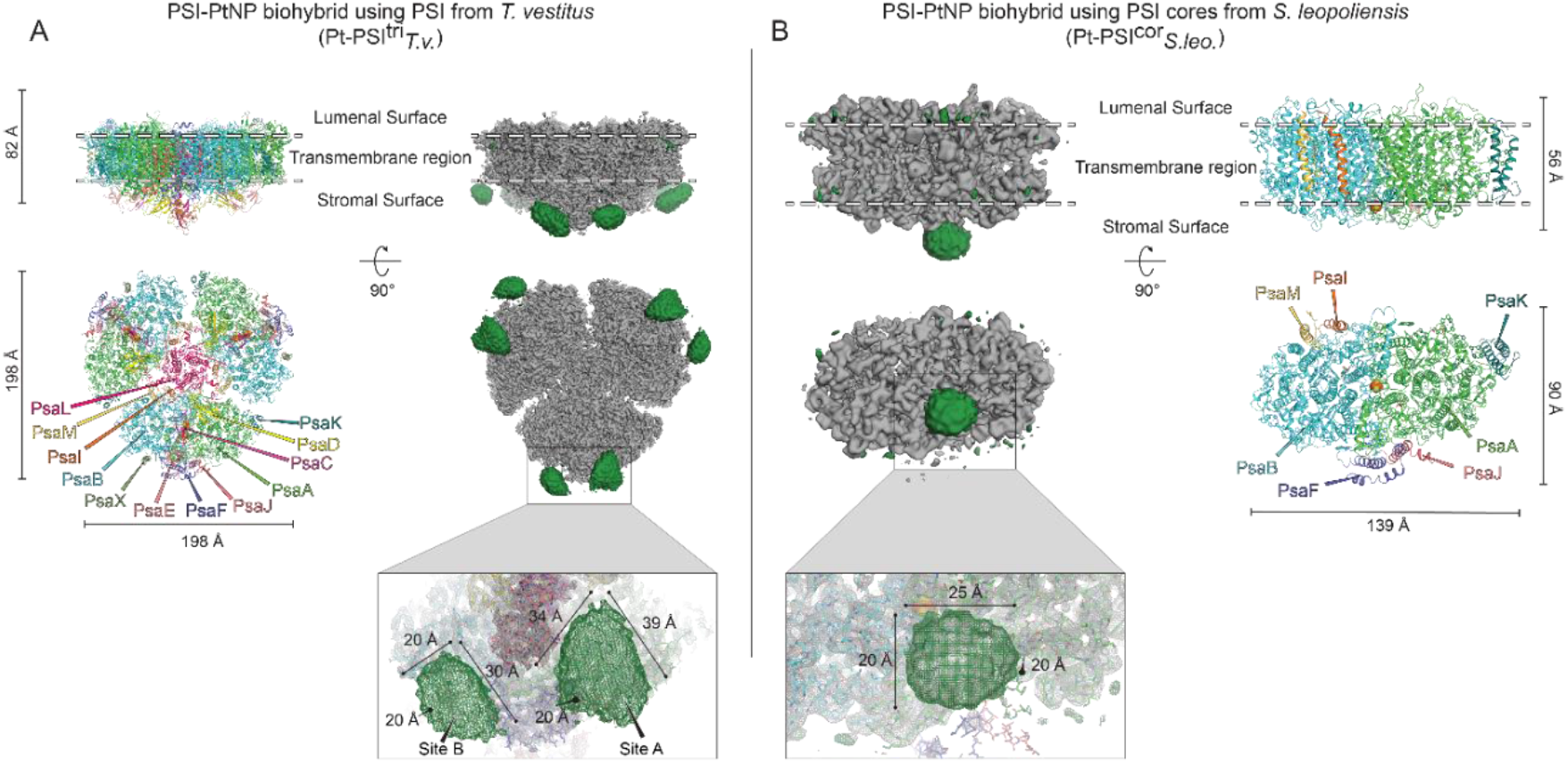
Pt-PSI^tri^_*T*.*v*._ and Pt-PSI^cor^_*S*.*leo*._ structures and PtNP signal measurements. (*A*) The Pt-PSI_tri*T*.*v*._ model (left) and map (right) with PSI signal in gray and PtNP signal in green. (*B*) The Pt-PSI^cor^ _*S*.*leo*._ model (right) and map (left) with PSI signal in gray and PtNP signal in green. Expanded panels below show the approximate dimensions of the PtNP signals.

It is also possible that the size distribution of PtNPs contributes to heterogeneity. The size distribution of PtNPs used to generate the PSI-PtNP biohybrids was centered at ∼2 nm (33). To determine whether PSI binds preferentially PtNPs of different sizes, we developed a procedure to assess the size distribution of PtNPs based on the cryo-EM micrograph images. This also yielded a size distribution centered at ∼2 nm (**Materials and Methods** and *SI Appendix*, **Fig. S9**), suggesting against preferential binding of specifically sized PtNPs in the biohybrid preparation procedure.

The Pt-PSI^cor^_*S*.*leo*._ structure was determined to 3.57 Å global resolution (*SI Appendix*, **Fig. S10**). Its data statistics and processing workflow are shown in *SI Appendix*, **Table S2** and **Fig. S11**. It is monomeric, containing subunits PsaA, PsaB, PsaF, PsaI, PsaJ, PsaK, and PsaM. Thus, compared to cyanobacterial PSI holocomplex structures, it lacks PsaC, PsaD, PsaE, and PsaL (and PsaX present in some organisms such as *T. vestitus*). Whereas the Pt-PSI^tri^_*T*.*v*._ structure has the typical dimensions of a PSI trimer, the smaller Pt-PSI^cor^_*S*.*leo*._ structure is ∼53 Å by 136 Å by 81 Å (**Fig. 1**). The Pt-PSI^cor^_*S*.*leo*._ subunits coordinate 86 chlorophyll *a* molecules, 1 chlorophyll *a′* molecule, 12 β-carotenes, 2 phylloquinones, 1 [4Fe-4S] cluster, and 3 lipids. A single PtNP binding site is present, in a position similar to the native electron acceptor-binding site, located on the stromal side of the complex on PsaA near its interface with PsaB, PsaF, and PsaJ (**Fig. 1** and **Fig. 2**). The signal is nearly spherical, ∼20 Å in each dimension. The closest distance between the PtNP and the nearest electron transfer cofactor, the [4Fe-4S] cluster F_X_, is ∼10 Å. Although this signal is more spherical than the PtNP signals from the trimeric biohybrid structures, it also does not show discrete signals for individual atoms in the PtNP, indicating heterogenous PtNP orientation/occupancy, but relatively higher positional homogeneity. Like the Pt-PSI^tri^_*T*.*v*._ data, the micrograph-based PtNP size distribution was also centered at ∼2 nm (*SI Appendix*, **Fig. S9**).

**Fig. 2.**
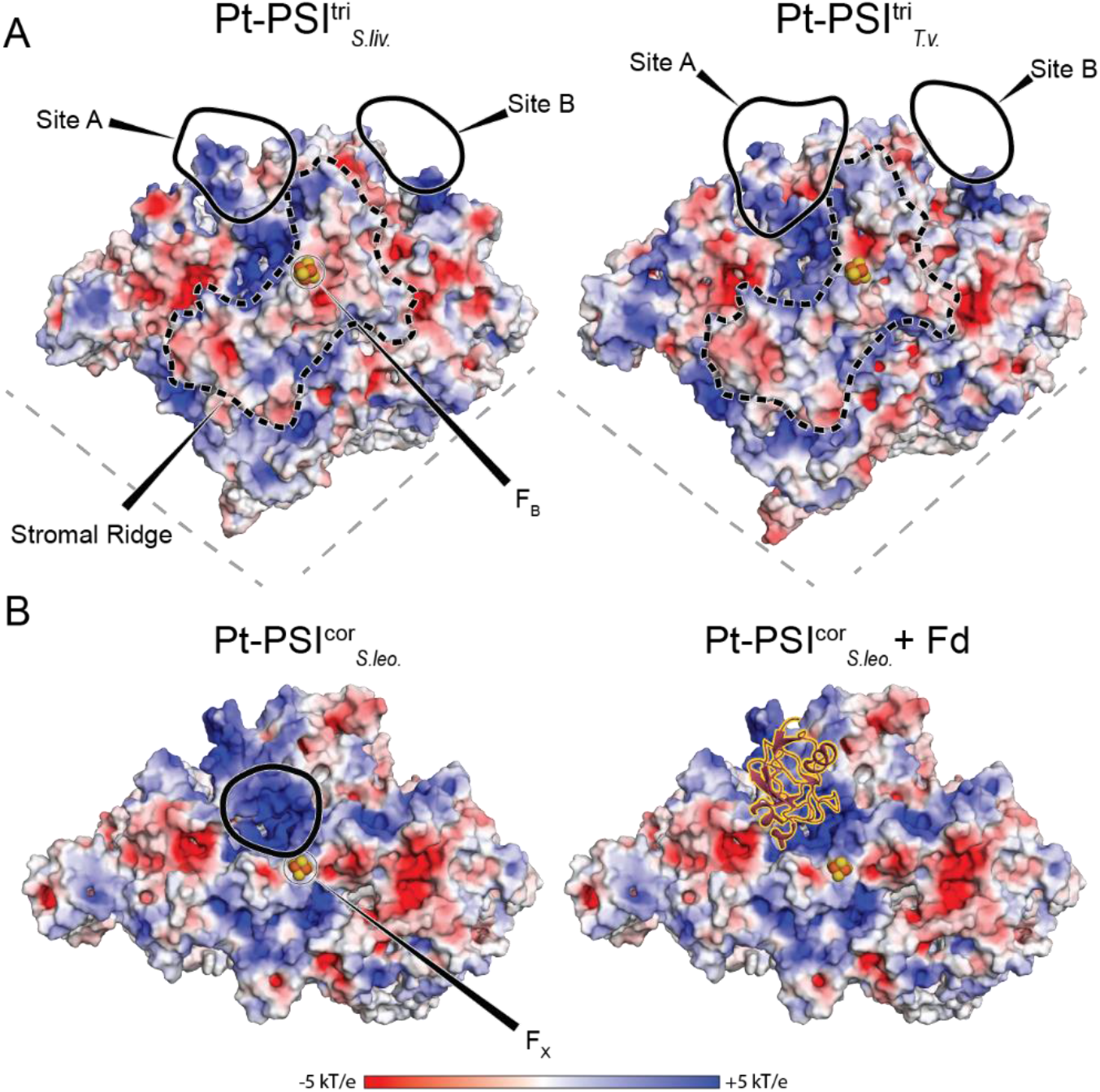
Removal of the stromal subunits allows PtNPs to bind closer to PSI electron transfer cofactors. (*A*) Electrostatic surfaces of Pt-PSI^tri^_*S*.*liv*._ (left) and Pt-PSI^tri^_*T*.*v*._ (right) structures. (*B*) Electrostatic surface of Pt-PSI^cor^_*S*.*leo*._ showing the PtNP location (left) relative to where Fd binds in the stromal ridge of trimeric PSI holocomplexes (right, based on PDB 7FIX). Solid black outlines show the locations of PtNP-binding sites and dashed black outlines show the region of the stromal ridge. For trimeric complexes, electrostatic surfaces were generated for only a single monomer for simplicity and light grey dashed lines denote the monomer-monomer interfaces. Electrostatic surfaces were calculated in PyMOL (41) using the APBS (42) plugin.

### Structure-based insight into PtNP-PSI binding interactions

Previous work suggested that electrostatic interactions are the primary driving force for PtNPs binding to PSI (20, 33). To investigate this, we compared electrostatic surfaces of the Pt-PSI^tri^_*T*.*v*._ structure, the Pt-PSI^tri^_*S*.*liv*._ structure that was reported previously, and the Pt-PSI^cor^_*S*.*leo*._ structure (**Fig. 2**). For the trimeric PSI structures, which are derived from different species of cyanobacteria, the electrostatic surfaces are highly similar. The primary positive region is surrounded by the stromal ridge subunits at the center of each monomer, near F_B_, forming the native acceptor binding site. The Pt-PSI^cor^_*S*.*leo*._ structure also shows a positively charged patch in this region (**Fig. 2**), highlighting the fact that core residues contribute greatly to the positive charge character of the native acceptor binding site in PSI holocomplexes. PtNP site A of the trimeric PSI structures overlaps slightly with the positively charged patch of the native acceptor binding site, but the PtNP site from the Pt-PSI^cor^_*S*.*leo*._ structure overlaps to a much greater extent, being positioned almost exactly where the native acceptors bind. This implies that the stromal subunits PsaC, PsaD, and PsaE actually hinder PtNPs from accessing the native acceptor binding site near the terminal [4Fe-4S] clusters, and their removal allows greater access. It also nicely shows that steric effects are a major contributor to native acceptor binding, and despite the PtNPs being a similar size to native acceptors, their shape does not compliment well the binding site and therefore cannot penetrate as deeply into that site compared to the native acceptors; ∼5.0-5.7 Å separates the terminal [4Fe-4S] cluster of PSI from the electron carrier cofactor in native acceptors (37–40, 43). A simplified schematic representing these features is shown in **Fig. 3**.

**Fig. 3.**
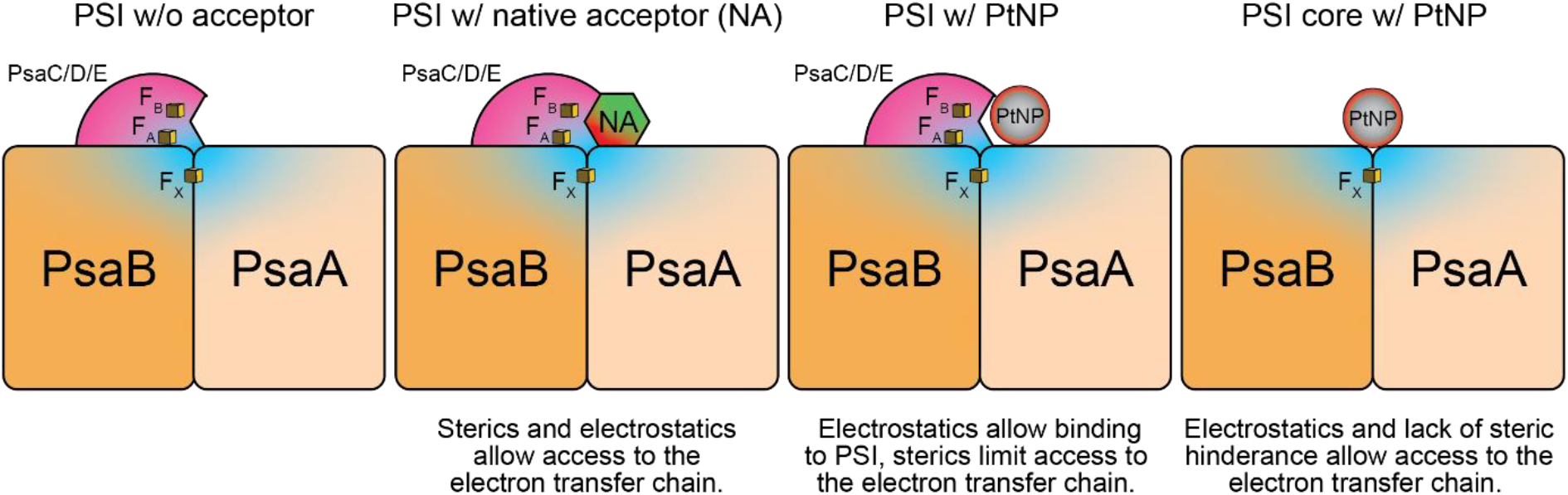
PSI core proteins bind electron acceptors electrostatically while steric effects limit access to the electron transfer chain. Only the terminal [4Fe-4S] clusters F_X_, F_A_, and F_B_ are shown, represented by yellow cubes. The blue and red coloring corresponds to positive and negative electrostatic surfaces, respectively.

There is much more ambiguity regarding whether charge interactions govern PtNP binding in site B of the trimeric structures. Although there is a clear positive patch at site B (**Fig. 2**), nearby residues toward the N-terminus of PsaX and the C-terminus of PsaE that were not resolved in the cryo-EM map and thus were not modelled in either the structure or MD simulations; however, these residues could interact with PtNPs (*SI Appendix*, **Fig. S12**). These regions are well conserved between *S. lividus* and *T. vestitus*, so if they do interact with the PtNP in site B, they probably do so similarly. It is noteworthy, however, that PSI from many cyanobacterial species lack PsaX, which could influence the binding of PtNPs in site B as considered further below.

### Molecular dynamics-based insight into PtNP-PSI binding interactions

Static cryo-EM structures poorly capture protein dynamics such as the impact of side chain rotamer states and loop dynamics on nanoscale interactions that vary over time. To obtain this information, we performed multiple 1 μs MD simulations on the Pt-PSI^tri^_*T*.*v*._ structure, the previously determined Pt-PSI^tri^_*S*.*liv*._ structure, and the Pt-PSI^cor^_*S*.*leo*._ structure. For each simulation, the PtNP diffusion coefficients are small, ∼2.2×10^−8^ cm^2^s^−1^ as measured by Einstein’s relation (*SI Appendix*, **Fig. S13**). This suggests that the PtNPs remain consistently associated with their binding site over the 1 μs trajectory.

To obtain insight into the residues prominently interacting with PtNPs, we performed a contact analysis based on the MD simulation (**Fig. 4** and *SI Appendix*, **Fig. S14**). Nearly all strong interactions arise from positively charged sidechains interacting with the mercaptosuccinic acid coating of the PtNP. In the Pt-PSI^tri^_*S*.*liv*._ and Pt-PSI^tri^_*T*.*v*._ structures, the interactions between the PtNP in site B and PSI arise primarily from residues in PsaB, especially Lys314, and additionally residues from PsaE, PsaF, and PsaX (**Fig. 4B** and *SI Appendix*, **Fig. S14**). Interestingly, the region containing PsaB-Lys314 is present in PSI from *T. vestitus* and *S. lividus*, both of which contain PsaX, but not PSI from *S. leopoliensis* or *Synechocystis* 6803, both of which lack PsaX (*SI Appendix*, **Fig. S5**). Thus, many of the interactions stabilizing the site B PtNP binding in *T. vestitus* and *S. lividus* may be absent in organisms lacking PsaX, and therefore the PSI from such organisms may not bind a PtNP in site B.

**Fig. 4.**
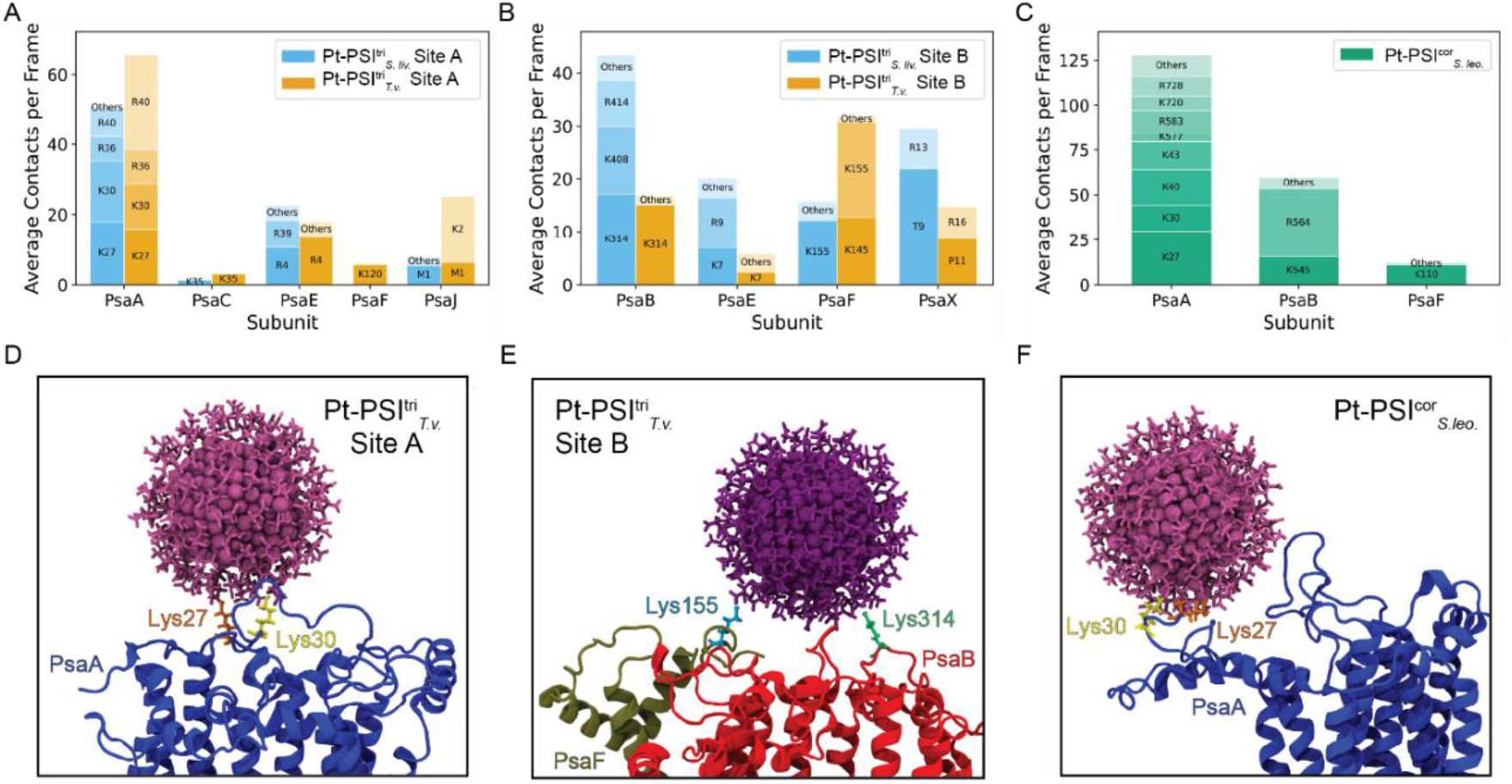
Molecular dynamics simulations show that positively charged residues dominate dynamic interactions. (*A*) Stacked contacts for PtNP site A in the Pt-PSI^tri^_*T*.*v*._ and Pt-PSI^tri^_*S*.*liv*._ structures. (*B*) Stacked contacts for PtNP site B in the Pt-PSI^tri^_*T*.*v*._ and Pt-PSI^tri^_*S*.*liv*._ structures. (*C*) Stacked contacts for the PtNP in the Pt-PSI^cor^_*S*.*leo*._ structure. (*D*) Single image of the MD simulation showing the contacts of PsaA-Lys 27 and Lys30 to the PtNP in site A of the Pt-PSI^tri^_*T*.*v*._ structure. (*E*) Same, but for PsaB-Lys 314 and PsaF-Lys155 to the PtNP in site B. (*F*) Same, but for PsaA-Lys 27 and Lys30 to the PtNP in the Pt-PSI^cor^ _*S*.*leo*._ structure.

The interactions between PSI and the PtNP in site A arise primarily from residues in PsaA (**Fig. 4A** and *SI Appendix*, **Fig. S14**). These comprise the sidechains of Lys27, Lys30, Arg36, and Lys/Arg40, and are additionally complemented by residues in PsaC, PsaE, PsaF, and PsaJ. PsaA-Lys27, Lys30, Arg36, and Lys/Arg40 and most of the other residues are conserved in *S. leopoliensis, T. vestitus, S. lividus*, and *Synechocystis* 6803 (*SI Appendix*, **Fig. S5**). This further suggests that the PSI from all four of these cyanobacteria, and probably many others, would bind a PtNP in site A similarly to what is observed in the Pt-PSI^tri^_*T*.*v*._ and Pt-PSI^tri^_*S*.*liv*._ structures.

In the Pt-PSI^cor^_*S*.*leo*._ structure, interactions between PSI and the PtNP are also dominated by positively charged sidechains from PsaA, especially the well conserved Lys27, Lys30, Arg36 residues that are also important for PtNP binding in site A of the Pt-PSI^tri^_*T*.*v*._ and Pt-PSI^tri^_*S*.*liv*._ structures (**Fig. 4** and *SI Appendix*, **Fig. S14**). Unlike the trimeric structures, however, the core lacks stromal ridge subunits (PsaC, PsaD, and PsaE), which allows the PtNP to interact with PsaB sidechains. The most prominent contacts are with the sidechains of Lys545 and Arg564 which are also well conserved among species (*SI Appendix*, **Fig. S5**). This suggests that PtNPs would bind in a similar location to PSI cores generated from other cyanobacterial species.

We further tested the likelihood of the most strongly interacting residues predicted by the contact analysis by calculating their minimum distance to their corresponding PtNPs (*SI Appendix*, **Fig. S15**). For the PtNP in site A of the Pt-PSI^tri^_*S*.*liv*._ and Pt-PSI^tri^_*T*.*v*._ structures, and the PtNP site in the Pt-PSI^cor^_*S*.*leo*._ structure, these are PsaA-Lys27 and Lys30, and for the PtNP in site B of the Pt-PSI^tri^_*S*.*liv*._ and Pt-PSI^tri^_*T*.*v*._ structures, these are PsaB-Lys314 and PsaF-Lys155. For the PtNP in site A of the trimeric structures, PsaA-Lys27 and Lys30 remain tightly associated for most of the 1 μs trajectory, with distances fluctuating around 1.5-2.0 Å, indicating persistent contact. Occasional transient spikes are observed corresponding to brief local rearrangements of the protein surface, but they do not disrupt the overall PtNP binding mode. The sustained proximity of PsaA-Lys27 and Lys30 supports their role as the primary stabilizing anchors that maintain PtNP binding at the PsaA interface.

Like the Pt-PSI^tri^_*S*.*liv*._ and Pt-PSI^tri^_*T*.*v*._ structures, the distance of PsaA-Lys27 to the PtNP in the Pt-PSI^cor^_*S*.*leo*._ structure remains strongly and consistently associated, ∼1.5-2.0 Å through the trajectory, with only rare deviations. PsaA-Lys30 also interacts robustly with the PtNP but exhibits greater fluctuation, including extended lengths of time (500 ns to 1 μs) where the distance increases to 3-6 Å. This suggests that while PsaA-Lys27 provides a persistent anchoring point for the PtNP, PsaA-Lys30 contributes a more dynamic interaction that modulates local binding flexibility without leading to full dissociation. The differences between PsaA-Lys27 and Lys30 interactions with the site A PtNP in trimeric structures and the PtNP site in the core structure are perhaps unsurprising because although these residues provide strong interactions in both cases, removal of the stromal ridge reveals additional residues of PsaA and PsaB that can also strongly interact with PtNPs (**Fig. 2** and **Fig. 4**).

In both the Pt-PSI^tri^_*S*.*liv*._ and Pt-PSI^tri^_*T*.*v*._ structures, PsaB-Lys314 maintains a short distance to the PtNP in site B, showing only occasional short-lived distance spikes, indicative of brief fluctuations rather than sustained dissociation. For PsaF-Lys155, the Pt-PSI^tri^_*S*.*liv*._ analysis shows intermittent periods of association and dissociation, including extended excursions beyond 4 to 8 Å, indicating dynamic but recurrent engagement with the PtNP, yet in the Pt-PSI^tri^_*T*.*v*._ analysis this residue is highly stable, with a distance remaining at ∼1.5–2.0 Å with only rare transient deviations. These PsaF-Lys155 differences could be interpreted as functionally relevant, but we emphasize caution due to previously described challenges to structural modelling in this region (*SI Appendix*, **Fig. S12**) that carry over to the MD simulations.

The initial contact-based analysis does not distinguish the identity of connecting atoms. Since we identify so many potential salt-bridges (**Fig. 4**), we quantify these interactions further using the frequency at which a hydrogen-bonding geometry occurs (<3.2 Å between heavy atoms and a donor-hydrogen-acceptor bond angle no greater than 30°) between PSI and the PtNP. We refer to this as an interaction analysis (**Fig. 5** and *SI Appendix*, **Tables S3-S5**), which allows comparison of the frequency and stability of PtNP-PSI interactions across replicas. We observe that some key residues involved in the PsaA interaction with the PtNP may interact with multiple mercaptosuccinic acid molecules simultaneously. Thus, rather than each amino acid contributing a single interaction to the mercaptosuccinic acid coating of the PtNP (corresponding to an interaction frequency of 100%), many residues exhibit interaction frequencies well above 100%, indicating the presence of multiple simultaneous interactions that together form strong tethers. If disruption of each contact is at least as strong as a hydrogen bond (∼1 to 1.2 kcal mol^−1^), the binding free energy between the PtNP and PSI is ∼−10 to −20 kcal mol^−1^. This corresponds to sub-nM dissociation constants which can be compared to native electron acceptors that bind weakly, with μM dissociation constants corresponding to standard binding free energies of ∼−7 to −9 kcal mol^−1^ (44). This difference is therefore a likely contributor to why electron transfer from PSI to native acceptors is blocked following PtNP binding (33).

**Fig. 5.**
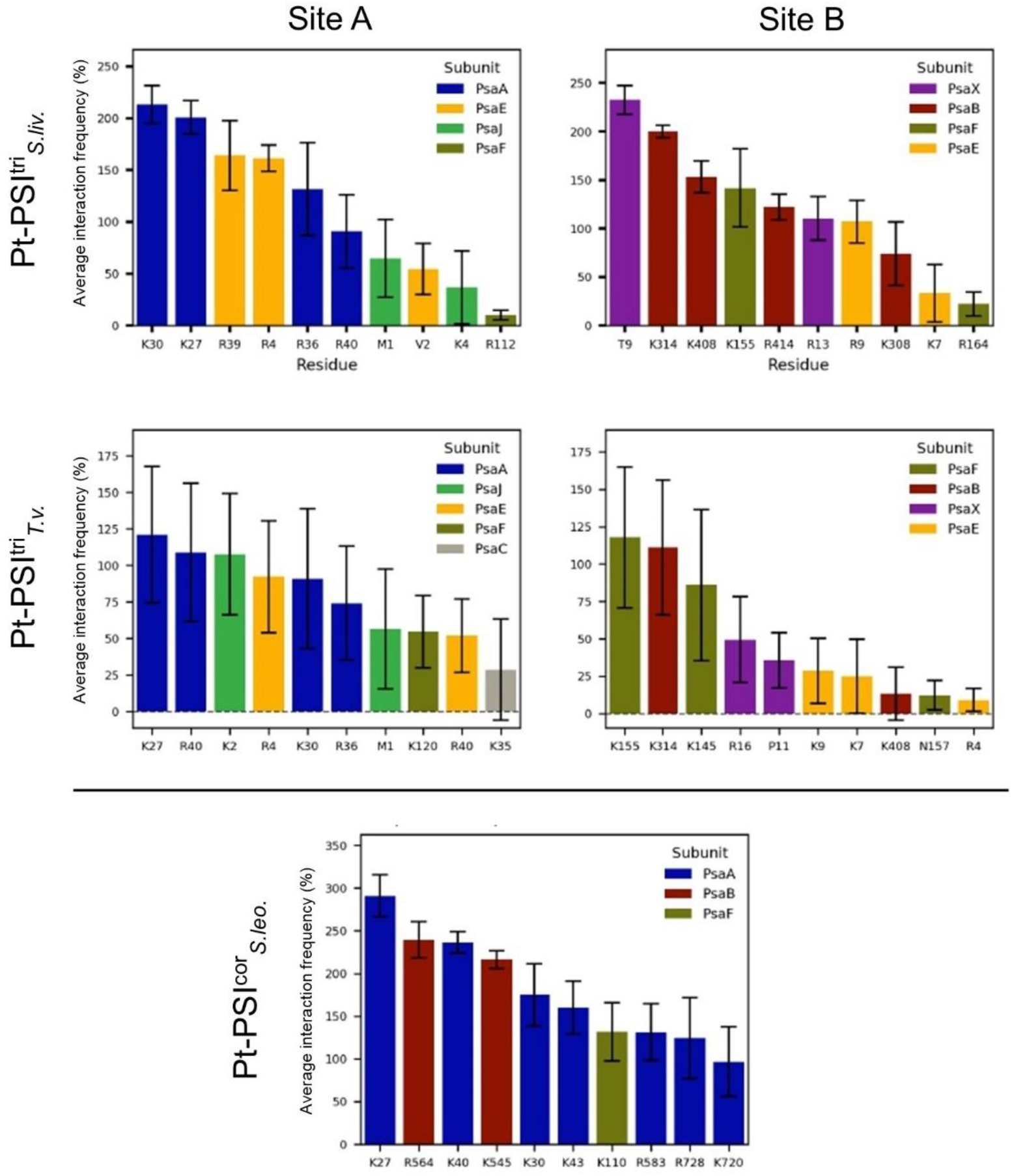
Interaction analysis highlights the most important residues to PtNP binding. The 10 residues contributing the most interactions between PSI and the bound PtNPs at each site are shown in order of their interaction frequency. Residues are color coded to their subunit’s identity. These interactions are tabulated in *SI Appendix*, **Table S3-S5**.

## Discussion

The ability to analyze a spheroidal abiotic electron acceptor (the PtNPs) provides unexpected insight into a fundamental feature of PSI: how it donates electrons to its native acceptors. Over billions of years, evolution has optimized the protein-protein binding interactions between PSI and its native acceptors. Canonically, that binding site comprises subunits PsaC, PsaD, and PsaE, with the terminal [4Fe-4S] clusters F_A_ and F_B_ coordinated by PsaC. Our MD simulations showed that for PtNP binding, the core PsaA subunit actually provides the most prominent electrostatic interactions rather than PsaC, PsaD, and PsaE (**Fig. 4** and **Fig. 5**). This seems likely to also be the case for native acceptors, as some of the positively charged residues involved in PtNP binding, such as PsaA-Arg36 and Arg/Lys40, additionally appear to participate in electrostatic interactions with native acceptors based on cryo-EM studies (37–40, 43).

Consistent with this, our electrostatic surface analysis showed that upon removal of PsaC, PsaD, and PsaE, PSI still maintains a positively charged patch onto which a PtNP can bind (**Fig. 2**). This is interestingly reminiscent of the heliobacterial reaction center, a PSI-like complex involved in anoxygenic photosynthesis (45, 46) thought to exhibit traits similar to the ancestor of extant reaction centers (47, 48). That reaction center lacks permanently bound stromal subunits and can promiscuously reduce soluble acceptors (46, 49) via their binding to a positive patch on the “P-side” of the protein (45, 50). Thus, it appears that PsaC, PsaD, and PsaE in PSI only partially provide complementary electrostatic interactions to native acceptors, and their more prominent role is for steering native acceptors via binding specificity that allow acceptor cofactors (e.g., a [2Fe-2S] cluster of Fd) to be positioned closely to the terminal PSI cofactor F_B_.

In the context of PtNP binding, PsaC, PsaD, and PsaE appear to inhibit close proximity to the electron transfer chain of PSI (**Fig. 3**). Originally, the mercaptosuccinic acid-coated PtNPs of ∼2 nm were selected to approximate the size and surface charge of native PSI electron acceptors, enabling electrostatic self-assembly of PSI-PtNP biohybrids. Although electron paramagnetic resonance spectroscopy and photocatalysis results demonstrated that PtNPs functionally mimic acceptor-protein binding to PSI (20, 31), the molecular details we have revealed delineate a heterogeneous binding landscape, especially in the productive site A of the biohybrids containing trimeric PSI. Namely, the oblong cryo-EM density of that site suggests that of the PtNPs binding in that vicinity, a relatively small population do so closely enough for efficient electron transfer (33). This is emphasized by our MD simulations showing that Pt atoms are rarely in close enough proximity to the terminal [4Fe-4S] cluster of PSI to participate in catalysis (*SI Appendix*, **Fig. S16**). This is consistent with the observation that under illumination, PtNP-PSI biohybrids containing trimeric PSI show H_2_ turnover frequencies of 1.5 to 1.8 H_2_ s^−1^ PSI^−1^, corresponding to an apparent electron delivery rate of 3.0 to 3.6 e^−^ s^−1^ per PSI (two electrons per H_2_), yet this is orders of magnitude slower than biological electron transfer rates expected, for example, across a 14 Å distance (10^5^ to 10^6^ s^−1^ in a protein matrix)(51). These observations suggest that only small PtNP subpopulations are likely to participate in H_2_ catalysis, highlighting the need for tighter control of PSI-PtNP biohybrids through enhanced manipulation strategies.

The donor-side electron supply to P_700_^+^ also seems likely to impose a kinetic bottleneck. *In vivo*, plastocyanin or cytochrome *c*_6_ (cyt *c*_6_) diffuse to a donor-side docking site (37, 52) to reduce P_700_^+^ before the next photochemical turnover. In our biohybrid system, although the intrinsic tunneling step from F_B_ to the PtNP is expected to be fast, productive H_2_ formation requires two successive light-generated electrons to reach the same PtNP. The second electron can only be delivered after P_700_^+^ has been reduced by cyt c_6_ (with sodium ascorbate as a sacrificial donor), a step that competes with charge recombination in PSI and thus suppresses the probability that both electrons reach the catalyst. This donor-side limitation has been repeatedly observed in PSI-hydrogenase and PSI-PtNP systems (15, 23, 25, 31).

In the core biohybrid, we removed the terminal [4Fe-4S] clusters F_A_ and F_B_, creating a PSI variant that can still drive H_2_ evolution via forward electron transfer, but now from the F_X_ cluster which is presumably more reducing. In this system, the distance between the PtNP and the terminal [4Fe-4S] cluster is shorter than that of the biohybrids containing trimeric PSI. For a typical protein medium, this decrease in donor-acceptor separation would be expected to strongly accelerate electron tunneling, and the higher driving force associated with electron transfer from F_X_ rather than F_B_ should further favor rapid forward transfer (an increase in driving force of ∼0.1 eV for F_X_ to Pt compared with F_B_ to Pt). Nonetheless, the observed H_2_ turnover frequency for Pt-PSI^cor^_*S*.*leo*._ is only 0.36 H_2_ s^−1^ PSI^−1^, corresponding to an apparent electron delivery rate of 0.72 e^−^ s^−1^ per PSI which is substantially slower than in the trimeric biohybrids. This striking result suggests that, in the Pt-PSI^cor^_*S*.*leo*._ system, H_2_ production is unlikely to be substantially limited by the intrinsic protein-catalyst tunneling step.

The reduced H_2_ activity of the core biohybrid underscores the dominant influence of donor-side limitations and recombination. Removal of F_A_ and F_B_ shortens the electron transfer chain of PSI, decreasing the lifetime of the charge-separated state from ∼65 ms (P_700_ ^+^F_B_ ^−^) to ∼1 ms (4, 34). This greatly increases competition between forward electron transfer from F_X_ ^−^ to the PtNP and unproductive charge recombination from F_X_^-^ to P_700_^+^. Thus, even though the core biohybrid delivers a more strongly reducing electron at a shorter distance to the Pt nanoparticle, the accelerated recombination kinetics and donor-side bottleneck likely outweigh these advantages and lead to lower H_2_ production. The PSI electron transfer chain, with its cofactors exhibiting stepped redox potentials and terminal [4Fe-4S] clusters, is evolutionarily optimized to generate long-lived charge separation that allows donor-side carriers to re-reduce P_700_^+^ and suppress recombination. Our results highlight that preserving this native-like, long-lived charge separation is as important as optimizing the protein-catalyst interface for efficiency.

Together, the PSI-PtNP biohybrids define a clear set of design principles for next-generation PSI-based solar fuel systems. First, solely shortening the electron-transfer pathway and increasing driving force at the acceptor side is insufficient to maximize H_2_ evolution; donor-side electron supply and suppression of charge recombination must be engineered in parallel. Strategies that provide rapid, sustained P_700_ ^+^ reduction—such as covalently tethered redox partners (15, 25), redox-polymer matrices (53, 54), or direct wiring of PSI to conductive electrode surfaces (55–58)—could relieve the donor-side bottleneck and allow the high intrinsic [4Fe-4S] to PtNP tunneling rates to be realized. Second, our structures show that interface geometry and electrostatics can be tuned to bias catalyst binding toward productive configurations. Specifically, exploiting the conserved, positively charged core surface near F_X_, or tailoring stromal-ridge topology and catalyst size/shape, offers routes to increase the fraction of PtNP sites positioned within effective tunneling distance of the terminal electron transfer cofactors in PSI. Finally, access to the more reducing F_X_-level electrons, once donor-side limitations are overcome, should enable PSI-based biohybrids to drive more challenging multi-electron transformations beyond H_2_ evolution, including CO_2_ reduction and other fuel-forming reactions. Thus, the structural and kinetic constraints identified here not only rationalize current performance limits but also point to concrete engineering pathways for optimizing PSI-nanomaterial architectures.

## Materials and Methods

### *T. vestitus* PSI isolation

*T. vestitus* cells were cultured at 45 °C under 70 μE white light. Cells were pelleted by centrifugation at 5,000×*g* for 20 min and resuspended in buffer A which contained 50 mM MES at pH 6.5 (HCl) with 15 mM CaCl_2_ and 10 mM MgCl_2_. Cells were lysed using a Siemens Microfluidics M-110P microfluidizer at 20,000 psi. Lysate was centrifuged at 6,000×*g* to remove unbroken cells. Thylakoid membranes were pelleted by ultracentrifugation (120,000×*g* at 4 °C for 30 min). Pelleted membranes were resuspended in buffer A at ∼0.5 mg chlorophyll mL^−1^. The membranes were solubilized by adding *n*-dodecyl-β-D-maltoside (β-DDM) to a concentration of 1% (w/v) while stirring in the dark for 1 h. Insoluble material was pelleted by centrifugation (25,000×*g* at 4 °C for 20 min). The supernatant was loaded onto 5-20% (w/v) sucrose gradients in buffer A with 0.01% β-DDM and ultracentrifuged (140,000×*g* at 4 °C for 18 h). The band containing PSI was collected, washed, and concentrated to ∼0.5 mg chlorophyll mL^−1^ with buffer A+0.03% β-DDM using an Amicon 50 kDa MWCO microfugal filter. The sample was then loaded onto another 5-20% (w/v) sucrose gradient and ultracentrifuged (140,000×*g* at 4 °C for 18 h). The band containing PSI trimers was collected and buffer exchanged into 20 mM Tricine at pH 8.0 (NaOH) with 75 mM NaCl and 0.05% β-DDM.

### *S. leopoliensis* PSI isolation and core generation

*S. leopoliensis* cells were grown in D_2_O (99.6%) AC (59) medium at 24 °C under 18 μE fluorescent lighting. Cells were washed and resuspended in 50 mM Tris-Cl pH 8.0 with 3 mM ethylenediaminetetraacetic acid. Cells were lysed with 0.1 mm glass beads in a pre-chilled Bead-Beater (BioSpec Products, Inc). The sample was beaten for eight 15-s bursts, with 4 min cooling in between runs in a surrounding ice bath. The solution was decanted and centrifuged at 907×*g* for 5 min in a Beckman Coulter Avanti JXN-26 centrifuge with a JLA 16.25 rotor at 4 °C. The supernatant was collected and loaded on top of a buffer containing 0.5 M sucrose, 10 mM ethylenediaminetetraacetic acid, and 50 mM Tris-Cl pH 8.0 (10 mL cell slurry: 15 mL buffer per tube). The thylakoid membranes were pelleted by ultracentrifugation for 1 h at 41,100×*g* in a Beckman L-60 ultracentrifuge with a 60 Ti rotor at 4 °C.

The sample was resuspended in 50 mM MES pH 6.6 with 10 mM MgCl_2_, and 15 mM CaCl_2_ to a concentration of 0.5 mg chlorophyll mL^−1^ and was solubilized with 1.0% β-DDM by gentle stirring for 1 h at 4 °C. Insoluble material was removed by centrifugation (JA 30.50 rotor, 14,100×*g*, 20 min). The supernatant was loaded onto 20 – 50% continuous sucrose density gradients prepared in 50 mM MES pH 6.6 with 10 mM MgCl_2_, 15 mM CaCl_2_, and 0.05% β-DDM. The middle portion of the dominant PSI trimer band was collected after ultracentrifugation (140,000×*g* for 17 h). The sucrose was removed by repeated dilution/concentration steps with Amicon Ultra-15 mL 50-kDa MWCO filtration devices and 50 mM MES pH 6.6 with 10 mM MgCl_2_, 15 mM CaCl_2_, and 0.02% β-DDM. The sample was concentrated to ∼3.0 mg chlorophyll mL^−1^ with a 100-kDa MWCO Amicon Ultra 0.5 mL spin concentrator.

To generate PSI cores (Pt-PSI^cor^_*S*.*leo*._), PSI at 0.9 mg chlorophyll mL^−1^ was incubated in a buffer containing 6.8 M urea, 50 mM Tris-Cl, and 75 mM glycine-NaOH pH 10.0 for 1 h as previously described (36). The PSI sample was then dialyzed overnight against 50 mM Tris-HCl pH 8.1. After dialysis, 0.02% β-DDM was added to the sample prior to concentration with 50-kDa MWCO Amicon Ultra-15 spin concentrators. Protein concentration was determined based on chlorophyll content in 100% methanol by measuring the absorbance at 665 nm. The sample, analyzed by inductively coupled plasma atomic emission spectroscopy on a ThermoScientific iCAP 6000 spectrometer, showed a ratio of ∼4 Fe/PSI monomer.

### Pt-PSI^cor^_*S*.*leo*._ and Pt-PSI^tri^_*T*.*v*._ biohybrid preparation

The synthesis of mercaptosuccinic acid-stabilized PtNPs was carried out according to literature procedures (60). Scanning transmission electron microscopy characterization confirmed the PtNPs to be crystalline, with a size distribution of 1.77±0.39 nm as previously reported (32) and is consistent with measurements taken from cryo-EM micrographs (*SI Appendix*, **Fig. S9**). Mercaptosuccinic acid-stabilized PtNPs (∼7.2 μM for diameter = 2.0 nm PtNPs) were added to the *S. leopoliensis* PSI (∼3 μM monomer) in a volume:volume ratio of 2:1 with an estimated 4.6 mol equivalents of PtNPs to one mol PSI monomer in a solution containing 50 mM Tris-Cl (pH 8.1) and 0.02% β-DDM and to *T. vestitus* PSI trimers in a volume:volume ratio of 5:1 with an estimated 9.0 mol equivalents of PtNPs to one mol PSI monomer in solution in buffer containing 20 mM Tris-Cl pH 8.0 and 0.05% β-DDM. The mixtures were tumbled overnight in the dark at 4 °C. Unbound PtNPs were removed from protein-bound nanoparticles by microfiltration (100-kDa MWCO Amicon Ultra-0.5 spin concentrators). The PSI-PtNP complexes were washed four times by repeated resuspension/concentration steps using microfiltration with buffer containing 20 mM Tris-Cl pH 8.1 and 0.02% β-DDM. Inductively coupled plasma atomic emission spectroscopy was used to determine the Pt and Fe content in the PSI-PtNP complexes. The PSI protein concentration was determined by Fe analysis assuming 4 Fe per PSI monomer. Based on this, an average of 344 ± 4 Pt atoms per PSI monomer was determined for the Pt-PSI^cor^_*S*.*leo*._ sample used for cryo-EM, and an average of 474 ± 5 Pt atoms per PSI monomer for the Pt-PSI^tri^_*T*.*v*._ sample. Based on the geometry-optimized PtNP model comprising 405 Pt atoms (33), approximately 0.85 PtNP are bound per PSI monomer in Pt-PSI^cor^_*S*.*leo*._ and 1.2 PtNP are bound per PSI monomer in Pt-PSI^tri^_*T*.*v*._.

### Photocatalytic hydrogen generation

Photocatalytic H_2_ production was initiated using a white light LED (Solis-3C, Thorlabs) and performed as previously described (31) utilizing 1.) 35 nM Pt-PSI^cor^_*S*.*leo*._ monomer and 12 μM cyt *c*_6_ or 2.) 25 nM Pt-PSI^tri^_*T*.*v*._ monomer and 10 μM cyt *c*_6_ with 10 mM MES pH 6.07, 100 mM sodium ascorbate, and 0.03% β-DDM in a total sample volume of 3.0 mL. An Agilent 990 micro gas chromatograph with a Molesieve 5A, 10 m/1 m column with a manual luer lock injector and UHP N_2_ carrier gas was used to detect H_2_ from samples of the headspace. The gas chromatograph instrument was calibrated using certified calibration H_2_ gas standards (Calibration Technologies Inc (CTI) and GasCo).

### Size exclusion chromatography

Size-exclusion chromatography was performed using a Superose 6 increase 5/150 GL (Cytiva) column equilibrated with 50 mM MES pH 6.5, 15 mM CaCl_2_, 10 mM MgCl_2_, 0.02% (w/v) β-DDM on a BioRad NGC Chromatography System. A 50 μL aliquot of PSI sample at 0.9 mg chlorophyll mL^−1^ concentration was injected into a 1 mL loading loop. The sample was then loaded onto the column, and the protein peaks were separated at a flow rate of 0.3 mL/min.

### Negative stain electron microscopy

For both biohybrids (Pt-PSI^tri^_*T*.*v*._ and Pt-PSI^cor^_*S*.*leo*._), 3 μL of sample at ∼1 μg chlorophyll mL^−1^ was applied to Formvar carbon film 400 mesh grids that were glow discharged at 20 mA for 7 s. The grids were incubated for one min and liquid was wicked away using filter paper. The grid was washed twice with 20 mM Tricine at pH 8.0 (NaOH) with 75 mM NaCl and 0.05% β-DDM and stained with uranyl acetate three times, incubating for one min each and wicking away liquid each time. Grids containing Pt-PSI^tri^_*T*.*v*._ biohybrids were imaged on a FEI CM120 transmission electron microscope operated at 80 kV with a BIOSPR12 camera. Grids containing Pt-PSI^cor^_*S*.*leo*._ biohybrids were imaged on a ThermoFisher F200C transmission electron microscope operated at 80 kV with a ThermoScientific Ceta 16M camera. Representative micrographs are shown in *SI Appendix*, **Fig. S4**.

### Cryo-EM grid preparation and data collection

Quantifoil 1.2/1.3 300 mesh cryo-EM grids were glow discharged for 30 s at 20 mA. Grids were mounted in a Thermo Fisher Vitrobot Mark IV system set to 4 °C and 100% humidity and 3 μL of either biohybrid at a concentration of ∼1 mg chlorophyll mL^−1^ was pipetted onto the grids. Grid were blotted (blot time=3 s), plunged into liquid ethane, and stored in liquid nitrogen. Cryo-EM screening and data acquisition were performed on a Talos Arctica transmission electron microscope operated at 200 kV using a Gatan K3 direct electron detector. The magnification was ×79,000 corresponding to a pixel size of 1.064 Å. The target defocus range was set to −1 to −2.5 μm. The total dose was 50 e^−^ Å^−2^ delivered over 4.13 s. 3,092 micrograph movies of Pt-PSI^tri^_*T*.*v*._ biohybrids and 2,572 micrograph movies of Pt-PSI^cor^_*S*.*leo*._ biohybrids were collected using EPU. Representative micrographs are shown in *SI Appendix*, **Fig. S8** and **Fig. S11**.

### Cryo-EM data processing

For both data sets, processing was performed in CryoSPARC v4 (61). Patch motion correction was used to correct, align, and dose-weight micrographs, and CTF Estimation (CTFFIND4) (62) was used to estimate the contrast transfer function. For the Pt-PSI^tri^_*T*.*v*._ sample, Blob Picker was used to select the initial particle set of 906,919 particles which were curated using Inspect Particle Picks and extracted. Two rounds of 2D classification (50 classes) yielded 124,600 accepted particles. These were further classified with an Ab-Initio reconstruction (8 classes) yielding 19,345 particles in the single accepted class. The associated volume was refined using the re-extracted, full-resolution particles in Homogeneous Refinement with C1 symmetry followed by Non-uniform Refinement with C3 symmetry. This yielded a final global resolution of 3.39 Å based on the Gold-standard Fourier shell correlation (0.143) cutoff criterion (63). Class images and processing workflows are shown in *SI Appendix*, **Fig. S8**. Resolution plots, local resolution map, and angular distribution are shown in *SI Appendix*, **Fig. S7**.

For the Pt-PSI^cor^_*S*.*leo*._ sample, Blob Picker was used to select 788,262 particles which were curated using Inspect Particle Picks and extracted. Two rounds of 2D classification (50 classes) yielded 380,050 accepted particles. These were further classified through an 8-class Ab-Initio reconstruction yielding 53,382 particles in a single accepted class. The associated volume was refined using the re-extracted, full-resolution particles in Homogeneous Refinement followed by Non-uniform Refinement both with C1 symmetry. This yielded a final global resolution of 3.57 Å based on the Gold-standard Fourier shell correlation (0.143) cutoff criterion (63). Class images and processing workflows are shown in *SI Appendix*, **Fig. S11**. Resolution plots, local resolution map, and angular distribution are shown in *SI Appendix*, **Fig. S10**.

### Model building

For the Pt-PSI^tri^_*T*.*v*._ structure, the starting model was the X-ray crystallography structure of PSI from *T. vestitus* (PDB 1JB0) (64), and for the Pt-PSI^cor^_*S*.*leo*._ structure, homology models of each subunits were created (SwissModel) (65) using the templates of the subunits from the X-ray crystallography structure of PSI from *T. vestitus* (PDB 1JB0) (64). In both cases, the model was fit into the map using UCSF ChimeraX (66). Manual fitting of subunits and cofactors was performed in Coot (67), and automated refinement was performed using phenix.real_space_refine (68) in the Phenix software suite (69).

### Micrograph-based PtNP size distribution determination

Ten defect-free micrographs were selected from each dataset. Micrographs were then imported into Matlab (R2024b) using the bio-formats imaging toolbox, inverted, and rescaled. Each image was segmented using a Gaussian smoothed image to locate PtNP signals and the centroid of each was extracted. From each centroid, a single line was plotted beyond the size of the segmented mask in the x direction, and the intensity of all traces were collected for analysis (*SI Appendix*, **Fig. S9**). Final object counts plotted are 7,080 and 5,188 for Pt-PSI^tri^_*T*.*v*._ and Pt-PSI^cor^_*S*.*leo*._ respectively. The full code is provided at https://git.doit.wisc.edu/cals/labs/gisriel/nanoparticle_analysis.

### Molecular dynamics

Each PSI-PtNP model from the three species was simulated with NAMD 3.0.1 using a GPU resident integrator (70) and the CHARMM36 force field without additional biases to track interactions and dynamics over an aggregate of 18 μs.

#### System Assembly

PSI monomers from *S. lividus* (PDB ID: 8VU3) (33), *T. vestitus*, and *S. leopoliensis* were built up into simulatable structures using VMD 1.9.4a57 (71). Unresolved residues were remodeled individually using SWISS-MODEL (65), using the existing cryo-EM structures as the template. Individual chains were combined using VMD 1.9.4a57 (71) and fed into PDB2PQR (72) to assign protonation states via PropKa (73) with an assumed pH of 7.0. The remodeled protein chains were reassembled into the full PSI complex, adding the cofactors extracted from the original structure. Isoprenoid tails were truncated to the point that they were unresolved in the cryo-EM data. Structurally resolved lipids were retained. [4Fe-4S] clusters and chlorophylls are also present and are built using existing topologies (74–76). The [4Fe-4S] charges were set to −2 and ligated to adjacent Cys residues to complete their coordination.

Based on the PtNP sizes estimated from the cryo-EM density, a 405 atom platinum nanoparticle was generated with the CHARMM-GUI nanomaterial modeler (33, 77). Each Pt atom is uncharged with a mass of 195.084 atomic mass units. Surface atoms further than 85 Å from the PtNP center were identified and functionalized with mercaptosuccinic acid ligands using CHARMM-compatible parameters generated by CGenFF (78). The mercaptosuccinic acid-capped PtNPs were solvated in a cubic water box with 0.15 M NaCl and equilibrated through short MD simulations. Equilibrated PtNPs were placed at the cryo-EM-identified binding sites on PSI. Rigid body fitting and merging were performed using VMD and TopoTools to generate the final system.

A cyanobacterial membrane was generated using CHARMM-GUI (79). The membrane model was expanded to a 3×3 array using TopoTools::replicatemol to increase the surface area for protein embedding. The equilibrated PSI-PtNP complex was then inserted into the bilayer. The resulting PSI-PtNP membrane system was subsequently solvated using the solvate package in VMD (71), which was the starting point for the simulation. Representative visualizations of biohybrid models used in this study are shown in *SI Appendix*, **Fig. S17**.

#### Simulations

MD simulations were performed with GPU-resident NAMD 3.0.1 and its immediate predecessors (70). Each system was energy minimized for 1,000 steps and equilibrated briefly to check for initial stability before production. Production equilibrium simulations were carried out in explicit solvent using the CHARMM36 force field for proteins (80), lipids (81), carbohydrates (82), and small molecules (78). The TIP3P water model was used (83), and a 12 Å cutoff was applied for nonbonded interactions. Long-range electrostatics were treated using the particle-mesh Ewald method with a 1.2 Å grid spacing (84). All simulations were performed in the NPT ensemble at 298 K using a Langevin thermostat (85) and barostat (86). The barostat decoupled the membrane-normal (z) axis from the membrane plane to maintain semi-isotropic pressure coupling. Hydrogen bond lengths were constrained using the SETTLE algorithm (87), but we found that the model we used for mercaptosuccinic acid/Pt interactions was unstable above a 1.8 fs timestep. All the systems were simulated to a length of 1 μs. To increase confidence in the results, 8 of these trajectories were run for the Pt-PSI^tri^_*S*.*liv*._ model, and 5 were run for the Pt-PSI^tri^_*T*.*v*._ and Pt-PSI^cor^_*S*.*leo*._ models.

#### Trajectory analysis

All MD trajectories were processed and analyzed using custom Python–VMD scripts developed specifically for this molecular system. We concretely calculate the root mean square deviation, diffusion coefficients, and multiple interaction metrics such as intermolecular contact, hydrogen bonding, and distances. Detailed descriptions for these processes are available in the supporting information. Numerical operations and data handling were performed using NumPy (88), while Matplotlib (89) was used to generate figures.

If your research involved human or animal participants, please identify the institutional review board and/or licensing committee that approved the experiments. Please also include a brief description of your informed consent procure if your experiments involved human participants.

## Data Availability

The Pt-PSI^tri^_*T*.*v*._ and Pt-PSI^cor^_*S*.*leo*._ coordinates and their associated cryo-EM maps have been deposited into the Protein Data Bank and Electron Microscopy Data Bank under PDB accession codes 10EG and 10KF, and EMDB accession codes EMD-75106 and EMD-75232, respectively. All input scripts to build and run molecular simulations are made publicly available on Zenodo| https://doi.org/10.5281/zenodo.18235787, together with the analysis scripts used to make the figures and plots in this article.

## Acknowledgments

We thank Dr. Jan Kern at LBNL for generously providing *T. vestitus* cells. Some of this work was performed in the Cryo-EM Research Center (CEMRC) in the Department of Biochemistry at the University of Wisconsin-Madison, and we thank their staff for support and assistance in cryo-EM data collection. We also thank the Center for Quantitative Cell Imaging for staff support and use of the Talos F200C microscope. The Gisriel group is a member of the SBGrid consortium and some of the analyses herein were performed using software compiled by SBGrid. Molecular dynamics simulations by the Vermaas group used the resources of Michigan State University’s Institute for Cyber-Enabled Research.

## References

1. D. A. Cherepanov, et al., Primary charge separation within the structurally symmetric tetrameric Chl2APAPBChl2B chlorophyll exciplex in photosystem I. J. Photochem. Photobiol. B 217, 112154 (2021).

2. N. Srinivasan, J. H. Golbeck, Protein–cofactor interactions in bioenergetic complexes: The role of the A1A and A1B phylloquinones in Photosystem I. Biochim. Biophys. Acta Bioenerg. 1787, 1057–1088 (2009).

3. P. Jordan, et al., Three-dimensional structure of cyanobacterial photosystem I at 2.5 Å resolution. Nature 411, 909–917 (2001).

4. K. Sauer, P. Mathis, S. Acker, J. A. Van best, Electron acceptors associated with P-700 in Triton solubilized Photosystem I particles from spinach chloroplasts. Biochim. Biophys. Acta Bioenerg. 503, 120–134 (1978).

5. B. Ke, R. E. Hansen, H. Beinert, Oxidation-Reduction Potentials of Bound Iron-Sulfur Proteins of Photosystem I. Proc. Nat’l. Acad. Sci. 70, 2941–2945 (1973).

6. L. M. Utschig, S. R. Soltau, D. M. Tiede, Light-driven hydrogen production from Photosystem I-catalyst hybrids. Curr. Opin. Chem. Biol. 25, 1–8 (2015).

7. E. Greenbaum, Platinized Chloroplasts: A Novel Photocatalytic Material. Science 230, 1373–1375 (1985).

8. I. J. Iwuchukwu, et al., Self-organized photosynthetic nanoparticle for cell-free hydrogen production. Nat. Nanotechnol. 5, 73–79 (2010).

9. I. J. Iwuchukwu, et al., Optimization of photosynthetic hydrogen yield from platinized photosystem I complexes using response surface methodology. Int. J. Hydrogen Energy 36, 11684–11692 (2011).

10. J. F. Millsaps, B. D. Bruce, J. W. Lee, E. Greenbaum, Nanoscale photosynthesis: photocatalytic production of hydrogen by platinized photosystem I reaction centers. Photochem. Photobiol. 73, 630–635 (2001).

11. E. Greenbaum, Interfacial photoreactions at the photosynthetic membrane interface: an upper limit for the number of platinum atoms required to form a hydrogen-evolving platinum metal catalyst. J. Phys. Chem. 92, 4571–4574 (1988).

12. J. W. Lee, C. V. Tevault, S. L. Blankinship, R. T. Collins, E. Greenbaum, Photosynthetic Water Splitting: In situ Photoprecipitation of Metallocatalysts for Photoevolution of Hydrogen and Oxygen. Energy Fuels 8, 770–773 (1994).

13. N. S. Ponomarenko, et al., Structural characterization of the platinum nanoparticle hydrogen-evolving catalyst assembled on Photosystem I by light-driven chemistry. ACS Nano 19, 4170–4185 (2025).

14. J. W. Lee, I. Lee, E. Greenbaum, Imaging nanometer metallocatalysts formed by photosynthetic deposition of water-soluble tansition metal compounds. J. Phys. Chem. B 109, 5409–5413 (2005).

15. C. E. Lubner, et al., Solar hydrogen-producing bionanodevice outperforms natural photosynthesis. Proc. Nat’l. Acad. Sci. 108, 20988–20991 (2011).

16. M. Gorka, J. H. Golbeck, Generating dihydrogen by tethering an [FeFe]hydrogenase via a molecular wire to the A1A/A1B sites of Photosystem I. Photosynth. Res. 143, 155–163 (2020).

17. C. E. Lubner, R. Grimme, D. A. Bryant, J. H. Golbeck, Wiring Photosystem I for Direct Solar Hydrogen Production. Biochemistry 49, 404–414 (2010).

18. A. Kanygin, et al., Rewiring photosynthesis: a photosystem I-hydrogenase chimera that makes H2 in vivo. Energy Environ. Sci. 13, 2903–2914 (2020).

19. I. Yacoby, et al., Photosynthetic electron partitioning between [FeFe]-hydrogenase and ferredoxin:NADP+-oxidoreductase (FNR) enzymes in vitro. Proc. Nat’l. Acad. Sci. 108, 9396–9401 (2011).

20. L. M. Utschig, et al., Photocatalytic Hydrogen Production from Noncovalent Biohybrid Photosystem I/Pt Nanoparticle Complexes. J. Phys. Chem. Lett. 2, 236–241 (2011).

21. R. A. Grimme, C. E. Lubner, J. H. Golbeck, Maximizing H2 production in Photosystem I/dithiol molecular wire/platinum nanoparticle bioconjugates. Dalton Trans 45, 10106–10113 (2009).

22. M. Miyachi, et al., A Photochemical Hydrogen Evolution System Combining Cyanobacterial Photosystem I and Platinum Nanoparticle-terminated Molecular Wires. Chem. Lett. 46, 1479–1481 (2017).

23. M. Gorka, J. Schartner, A. van der Est, M. Rögner, J. H. Golbeck, Light-Mediated Hydrogen Generation in Photosystem I: Attachment of a Naphthoquinone–Molecular Wire–Pt Nanoparticle to the A1A and A1B Sites. Biochemistry 53, 2295–2306 (2014).

24. K. A. Walters, J. H. Golbeck, Designing a modified clostridial 2[4Fe-4S] ferredoxin as a redox coupler to directly link photosystem I with a Pt nanoparticle. Photosynth. Res. 143, 165–181 (2020).

25. B. R. Evans, H. M. O’Neill, S. A. Hutchens, B. D. Bruce, E. Greenbaum, Enhanced Photocatalytic Hydrogen Evolution by Covalent Attachment of Plastocyanin to Photosystem I. Nano Lett. 4, 1815–1819 (2004).

26. R. A. Grimme, C. E. Lubner, D. A. Bryant, J. H. Golbeck, Photosystem I/molecular wire/metal nanoparticle bioconjugates for the photocatalytic production of H2. J. Am. Chem. Soc. 130, 6308–6309 (2008).

27. M. Gorka, et al., Electron transfer from the A1A and A1B sites to a tethered Pt nanoparticle requires the FeS clusters for suppression of the recombination channel. J. Photochem. Photobiol. B 152, 325–334 (2015).

28. S. C. Silver, et al., Protein delivery of a Ni catalyst to Photosystem I for light-driven hydrogen production. J. Am. Chem. Soc. 135, 13246–13249 (2013).

29. L. M. Utschig, S. C. Silver, K. L. Mulfort, D. M. Tiede, Nature-driven photochemistry for catalytic solar hydrogen production: a Photosystem I-transition metal catalyst hybrid. J. Am. Chem. Soc. 133, 16334–16337 (2011).

30. A. H. Teodor, et al., Aqueous-soluble bipyridine cobalt(II/III) complexes act as direct redox mediators in photosystem I-based biophotovoltaic devices. RSC Adv. 11, 10434–10450 (2021).

31. L. M. Utschig, S. R. Soltau, K. L. Mulfort, J. Niklas, O. G. Poluektov, Z-scheme solar water splitting via self-assembly of photosystem I-catalyst hybrids in thylakoid membranes. Chem. Sci. 9, 8504–8512 (2018).

32. L. M. Utschig, N. J. Zaluzec, T. Malavath, N. S. Ponomarenko, D. M. Tiede, Solar water splitting Pt-nanoparticle photosystem I thylakoid systems: Catalyst identification, location and oligomeric structure. Biochim. Biophys. Acta Bioenerg. 1864, 148974 (2023).

33. C. J. Gisriel, et al., Structure of a biohybrid photosystem I-platinum nanoparticle solar fuel catalyst. Nat. Commun. 15, 9519 (2024).

34. M. Gorka, et al., Electron transfer from the A1A and A1B sites to a tethered Pt nanoparticle requires the FeS clusters for suppression of the recombination channel. J. Photochem. Photobiol. B 152, 325–334 (2015).

35. L.-O. Pålsson, J. P. Dekker, E. Schlodder, R. Monshouwer, R. van Grondelle, Polarized site-selective fluorescence spectroscopy of the long-wavelength emitting chlorophylls in isolated Photosystem I particles of Synechococcus elongatus. Photosynth. Res. 48, 239–246 (1996).

36. P. V. Warren, K. G. Parrett, J. T. Warden, J. H. Golbeck, Characterization of a photosystem I core containing P700 and intermediate electron acceptor A1. Biochemistry 29, 6545–6550 (1990).

37. J. Li, et al., Structure of cyanobacterial photosystem I complexed with ferredoxin at 1.97 Å resolution. Commun. Biol. 5, 951 (2022).

38. H. Kubota-Kawai, et al., X-ray structure of an asymmetrical trimeric ferredoxin– photosystem I complex. Nat. Plants 4, 218–224 (2018).

39. C. J. Gisriel, et al., Structure of a photosystem I-ferredoxin complex from a marine cyanobacterium provides insights into far-red light photoacclimation. J. Biol. Chem. 298, 101408 (2022).

40. P. Cao, et al., Structural basis for energy and electron transfer of the photosystem I-IsiA-flavodoxin supercomplex. Nat Plants 6, 167–176 (2020).

41. W. L. DeLano, The pyMol molecular graphics system.

42. E. Jurrus, et al., Improvements to the APBS biomolecular solvation software suite. Protein Sci. 27, 112–128 (2018).

43. I. Caspy, A. Borovikova-Sheinker, D. Klaiman, Y. Shkolnisky, N. Nelson, The structure of a triple complex of plant photosystem I with ferredoxin and plastocyanin. Nat. Plants 6, 1300–1305 (2020).

44. P. Q. Y. Setif, H. Bottin, Laser flash absorption spectroscopy study of ferredoxin reduction by Photosystem I: Spectral and kinetic evidence for the existence of several Photosystem I-ferredoxin complexes. Biochemistry 34, 9059–9070 (1995).

45. C. Gisriel, et al., Structure of a symmetric photosynthetic reaction center-photosystem. Science 357, 1021–1025 (2017).

46. B. Ferlez, et al., Thermodynamics of the Electron Acceptors in Heliobacterium modesticaldum: An Exemplar of an Early Homodimeric Type I Photosynthetic Reaction Center. Biochemistry 55, 2358–2370 (2016).

47. W. M. Sattley, et al., The Genome of Heliobacterium modesticaldum, a Phototrophic Representative of the Firmicutes Containing the Simplest Photosynthetic Apparatus. J. Bacteriol. 190, 4687–4696 (2008).

48. G. S. Orf, C. Gisriel, K. E. Redding, Evolution of photosynthetic reaction centers: insights from the structure of the heliobacterial reaction center. Photosynth. Res. 138, 11–37 (2018).

49. S. P. Romberger, J. H. Golbeck, The FX iron–sulfur cluster serves as the terminal bound electron acceptor in heliobacterial reaction centers. Photosynth. Res. 111, 285–290 (2012).

50. G. S. Orf, C. Gisriel, K. E. Redding, Evolution of photosynthetic reaction centers: insights from the structure of the heliobacterial reaction center. Photosynth. Res. 138, 11–37 (2018).

51. C. C. Moser, J. M. Keske, K. Warncke, R. S. Farid, P. L. Dutton, Nature of biological electron transfer. Nature 355, 796–802 (1992).

52. I. Caspy, A. Borovikova-Sheinker, D. Klaiman, Y. Shkolnisky, N. Nelson, The structure of a triple complex of plant photosystem I with ferredoxin and plastocyanin. Nat. Plants 6, 1300–1305 (2020).

53. A. Badura, et al., Photocurrent generation by photosystem 1 integrated in crosslinked redox hydrogels. Energy Environ. Sci. 4, 2435–2440 (2011).

54. T. Kothe, et al., Engineered electron-transfer chain in photosystem 1 based photocathodes outperforms Electron-Transfer Rates in natural photosynthesis. Chem. Eur. J. 20, 11029–11034 (2014).

55. O. Yehezkeli, et al., Integrated photosystem II-based photo-bioelectrochemical cells. Nat. Commun. 3, 742 (2012).

56. G. LeBlanc, G. Chen, G. K. Jennings, D. E. Cliffel, Photoreduction of Catalytic Platinum Particles Using Immobilized Multilayers of Photosystem I. Langmuir 28, 7952–7956 (2012).

57. K. Nguyen, B. D. Bruce, Growing green electricity: Progress and strategies for use of Photosystem I for sustainable photovoltaic energy conversion. Biochim. Biophys. Acta Bioenerg. 1837, 1553–1566 (2014).

58. K. R. Stieger, S. C. Feifel, H. Lokstein, F. Lisdat, Advanced unidirectional photocurrent generation via cytochrome c as reaction partner for directed assembly of photosystem I. Phys. Chem. Chem. Phys. 16, 15667–15674 (2014).

59. H. L. Crespi, S. M. Conrad, R. A. Uphaus, J. J. Katz, Cultivation of Microorganisms in Heavy Water. Ann. N. Y. Acad. Sci. 84, 648–666 (1960).

60. S. Chen, K. Kimura, Synthesis of Thiolate-Stabilized Platinum Nanoparticles in Protolytic Solvents as Isolable Colloids. J. Phys. Chem. B 105, 5397–5403 (2001).

61. A. Punjani, J. L. Rubinstein, D. J. Fleet, M. A. Brubaker, cryoSPARC: algorithms for rapid unsupervised cryo-EM structure determination. Nat. Methods 14, 290–296 (2017).

62. A. Rohou, N. Grigorieff, CTFFIND4: Fast and accurate defocus estimation from electron micrographs. J. Struct. Bio. 192, 216–221 (2015).

63. S. H. W. Scheres, S. Chen, Prevention of overfitting in cryo-EM structure determination. Nat. Methods 9, 853–854 (2012).

64. P. Jordan, et al., Three-dimensional structure of cyanobacterial photosystem I at 2.5 Å resolution. Nature 411, 909–917 (2001).

65. A. Waterhouse, et al., SWISS-MODEL: homology modelling of protein structures and complexes. Nucleic Acids Res. 46, W296–W303 (2018).

66. E. C. Meng, et al., UCSF ChimeraX: Tools for structure building and analysis. Protein Sci. 32, e4792 (2023).

67. P. Emsley, B. Lohkamp, W. G. Scott, K. Cowtan, Features and development of Coot. Acta Crystallogr. D Struct. Biol. 66, 486–501 (2010).

68. P. V. Afonine, et al., Real-space refinement in PHENIX for cryo-EM and crystallography. Acta Crystallogr. D Struct. Biol. 74, 531–544. (2018)

69. P. D. Adams, et al., PHENIX: a comprehensive Python-based system for macromolecular structure solution. Acta Crystallogr. D Struct. Biol. 66, 213–221 (2010).

70. J. C. Phillips, et al., Scalable molecular dynamics on CPU and GPU architectures with NAMD. J. Chem. Phys. 153, 044130 (2020).

71. W. Humphrey, A. Dalke, K. Schulten, VMD: visual molecular dynamics. J. Mol. Graph. 14, 33–38, 27–28 (1996).

72. T. J. Dolinsky, J. E. Nielsen, J. A. McCammon, N. A. Baker, PDB2PQR: an automated pipeline for the setup of Poisson-Boltzmann electrostatics calculations. Nucleic Acids Res. 32, W665–667 (2004).

73. T. J. Dolinsky, et al., PDB2PQR: expanding and upgrading automated preparation of biomolecular structures for molecular simulations. Nucleic Acids Res. 35, W522–W525 (2007).

74. M.-L. Tan, B. S. Perrin Jr., S. Niu, Q. Huang, T. Ichiye, Protein dynamics and the all-ferrous [Fe4S4] cluster in the nitrogenase iron protein. Protein Sci. 25, 12–18 (2016).

75. F. Guerra, S. Adam, A.-N. Bondar, Revised force-field parameters for chlorophyll-a, pheophytin-a and plastoquinone-9. J. Mol. Graph. 58, 30–39 (2015).

76. A. Damjanović, I. Kosztin, U. Kleinekathöfer, K. Schulten, Excitons in a photosynthetic light-harvesting system: A combined molecular dynamics, quantum chemistry, and polaron model study. Phys. Rev. E 65, 031919 (2002).

77. Y. K. Choi, et al., CHARMM-GUI Nanomaterial Modeler for Modeling and Simulation of Nanomaterial Systems. J. Chem. Theory Comput. 18, 479–493 (2022).

78. K. Vanommeslaeghe, et al., CHARMM general force field: A force field for drug-like molecules compatible with the CHARMM all-atom additive biological force fields. J. Comput. Chem. 31, 671–690 (2010).

79. S. Jo, T. Kim, V. G. Iyer, W. Im, CHARMM-GUI: A web-based graphical user interface for CHARMM. J. Comput. Chem. 29, 1859–1865 (2008).

80. J. Huang, et al., CHARMM36m: an improved force field for folded and intrinsically disordered proteins. Nat. Methods 14, 71–73 (2017).

81. J. B. Klauda, et al., Update of the CHARMM All-Atom Additive Force Field for Lipids: Validation on Six Lipid Types J. Phys. Chem. B 114, 7830–7843 (2010)

82. O. Guvench, A.D. MacKerell Jr. Comparison of protein force fields for molecular dynamics simulations. Methods Mol. Biol. 443, 63–88 (2008)

83. W. L. Jorgensen, J. Chandrasekhar, J. D. Madura, R. W. Impey, M. L. Klein, Comparison of simple potential functions for simulating liquid water. J. Chem. Phys. 79, 926–935 (1983).

84. U. Essmann, et al., A smooth particle mesh Ewald method. J. Chem. Phys. 103, 8577–8593 (1995).

85. R. Kubo, The fluctuation-dissipation theorem. Rep. Prog. Phys. 29, 255 (1966).

86. S. E. Feller, Y. Zhang, R. W. Pastor, B. R. Brooks, Constant pressure molecular dynamics simulation: The Langevin piston method. J. Chem. Phys. 103, 4613–4621 (1995).

87. S. Miyamoto, P. A. Kollman, Settle: An analytical version of the SHAKE and RATTLE algorithm for rigid water models. J. Comput. Chem. 13, 952–962 (1992).

88. C. R. Harris, et al., Array programming with NumPy. Nature 585, 357–362 (2020).

89. J. D. Hunter, Matplotlib: A 2D Graphics Environment. Comput. Sci. Eng. 9, 90–95 (2007).

