## Supplemental Information for "Molecular design principles for Photosystem I-based biohybrid solar fuel catalysts"

**This PDF file includes:**

Supporting text S1
Figs. S1 to S17
Table S1 to S5
Data S1
SI References

**Text S1. Extended Simulation Analysis Description.**
*RMSD and diffusion coefficient calculations for PtNPs*

To track PtNP stability over time, we measure the RMSD for the platinum atoms within the PtNP with respect to its initially modeled structure, and found that it was always less than 1 Å. Similarly, the RMSD for the PSI models themselves were tracked, and were found to be similar in magnitude to the cryo-EM resolution, as expected(1). Diffusion coefficients for each nanoparticle were determined by measuring the mean squared displacement for the PtNP center and then using Einstein's relation to measure diffusion(2). Concretely, we measured the diffusion for each PtNP in 9 ns chunks along the 1 μs trajectory. The mean squared displacement in three dimensions is related to the diffusion coefficient by:

$$D = \frac{MSD}{6\Delta t}$$

where MSD stands for mean squared displacement. Differences in diffusion behavior across replicas were quantified using the standard deviation of the diffusion coefficients derived from each independent simulation replica.

*Interactions between PSI and platinum nanoparticles*

The first of two approaches to quantify PSI-PtNP interactions is through a weighted residue-PtNP contact analysis. For each frame, all heavy atoms of PSI were evaluated against all PtNP atoms within a 6 Å search radius. To avoid a binary cutoff and better capture transient interactions, a smooth sigmoidal weighting function was used based on prior work(3):

$$Cij = 1/\left(1 + exp\left(5\left(d_{ij} - 4\text{Å}\right)\right)\right)$$

where $d_{ij}$ is the distance between PSI atom (i) and PtNP atom (j).

This approach produces fractional contact values that gradually decrease as distances exceed 4 Å. Weighted contributions were summed on a per-residue basis for each frame, and contact profiles were averaged across simulation replicas to identify residues involved in persistent PtNP interactions.

In addition to contact counting, minimum-distance measurements were performed to evaluate the proximity of specific PSI residues to the PtNP. For each frame, all heavy-atom distances between a selected residue and the PtNP were calculated, and the smallest value was recorded. This provided a direct indicator of whether a given residue remained closely associated with the PtNP during the simulation. Distance measurements were computed for each replica to assess consistency of residue-PtNP interactions.

*Interaction analysis between PSI and PtNPs*

To compare the frequency and stability of PtNP-PSI interactions across replicas, we leveraged the VMD h-bonds plugin as reported previously(4). For each replica, interactions were evaluated between all protein atoms and the interaction across the trajectories. A donor-acceptor distance cutoff of 3.2 Å(5) and a donor bond angle cutoff of 30°(4) were used to identify interactions. Interaction counts were computed for each frame and recorded separately for each PtNP, allowing comparison of the frequency and stability of PtNP-PSI interactions across replicas.

*Electron transfer distance measurements between platinum nanoparticles and [4Fe-4S] cluster*

To evaluate the potential for electron transfer between PSI and the PtNPs, we measured the shortest distance between the terminal [4Fe-4S] cluster in PSI and each PtNP throughout the simulations. We use an expansive definition for the [4Fe-4S] cluster, including the iron atoms, as well as all eight sulfur atoms that coordinate these irons (4 from the cluster, and 4 from the coordinating cysteine residues). For every trajectory frame, we computed the minimum distance between any of these atoms and the PtNP. Although minimum distances were computed between the PtNP and all potential interacting atoms, in practice the shortest contact was consistently contributed by a single cysteine sulfur: PsaC-Cys14 (Pt-PSI$^{tri}_{S.liv.}$), PsaC-Cys14 (Pt-PSI$^{tri}_{T.v.}$), and PsaA-Cys586 (Pt-PSI$^{cor}_{S.leo.}$). These time-dependent distances indicate how closely each PtNP remains positioned relative to the terminal [4Fe-4S] cluster, which is a key requirement for efficient electron transfer.

**Supplementary figures**

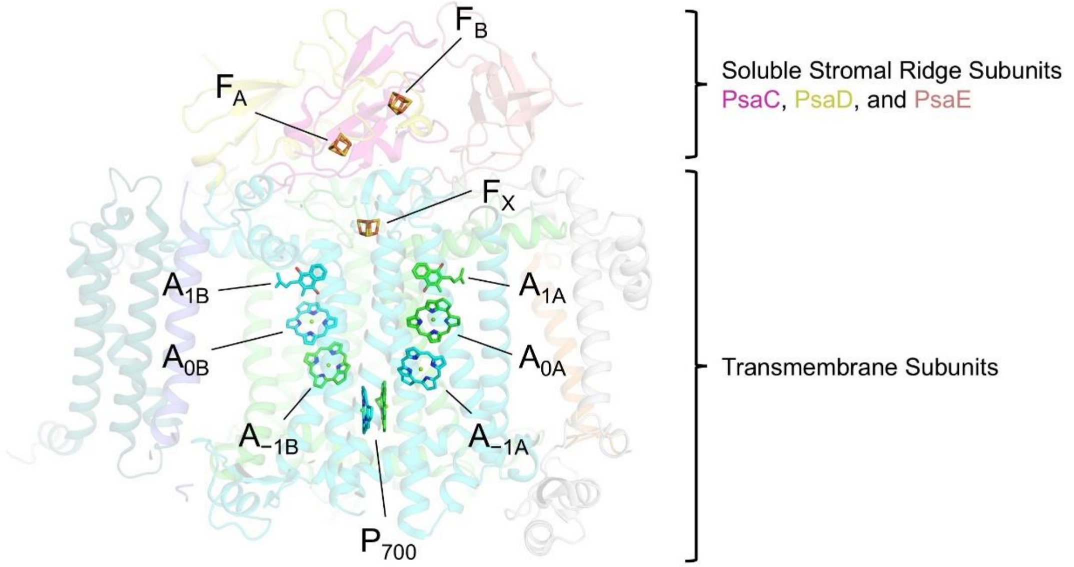


**Fig. S1.** PSI electron transfer chain and domains. Chlorophyll substituents and cofactor tails are hidden for clarity.

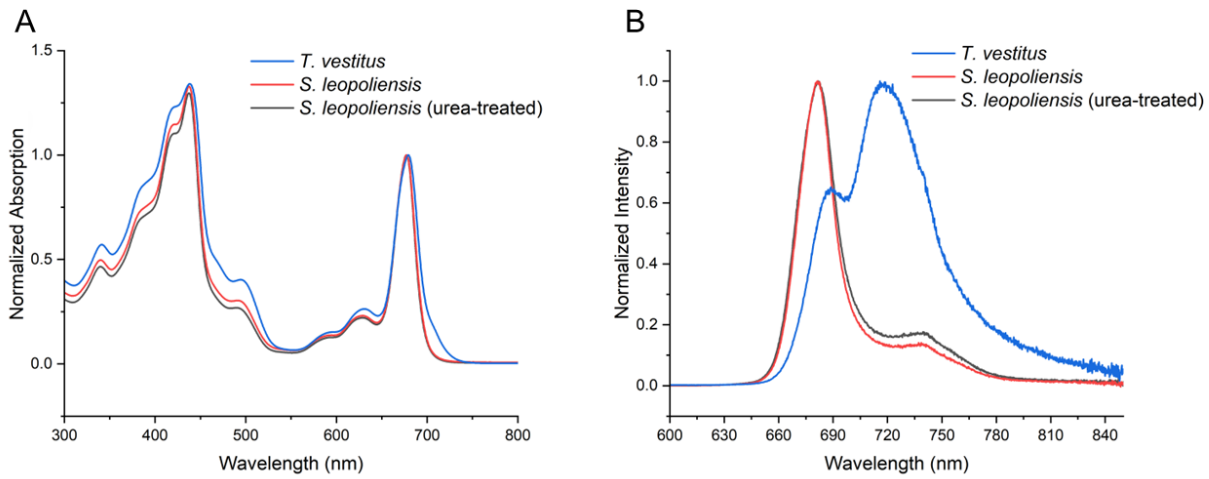


**Fig. S2.** Room temperature absorption and fluorescence emission spectra of PSI samples. (*A*) Room temperature absorption. (*B*) Room temperature fluorescence emission (ex. 440 nm).

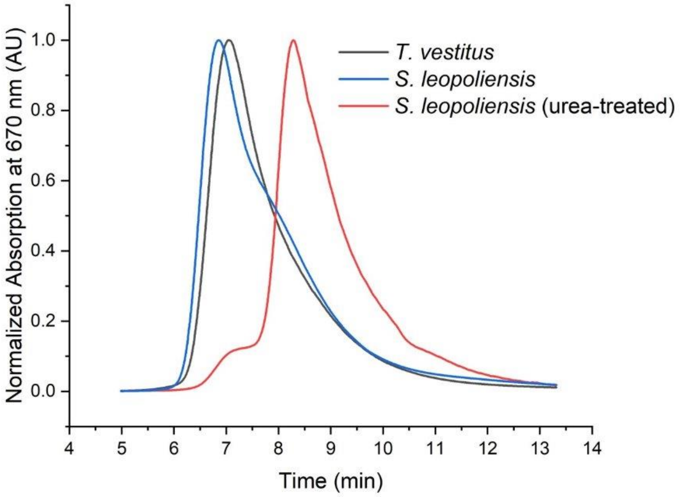


**Fig. S3.** Size exclusion chromatography of PSI samples. Absorbance was monitored at 670 nm.

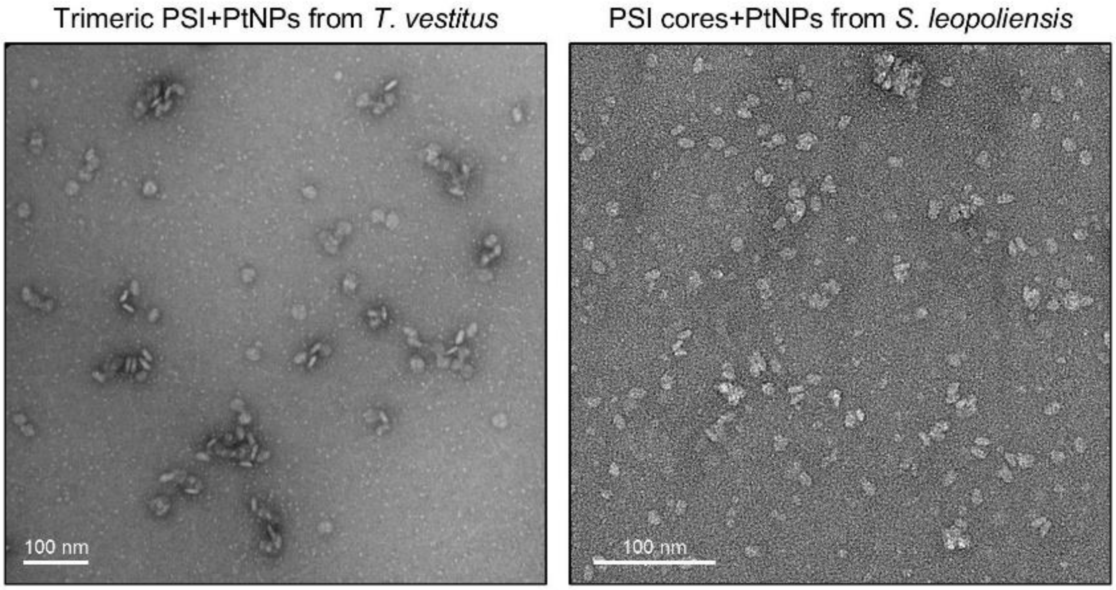


**Fig. S4.** Negative staining of biohybrids generated from *T. vestitus* trimers and *S. leopoliensis* cores.

### PsaA

```
S. leopoliensis    MTISPPEREAKVKATVDKNPVPTSFEKWGKPGHFDRTLAKGPKTTTWIWNLHANAHDFDS 60
T. vestitus        MTISPPEREPKVRVVVDNDPVPTSFEKWAKPGHFDRTLARGPQTTTWIWNLHALAHDFDT 60
S. lividus         MTISPPEREPKVRVVVDNDPVPTSFEKWAKPGHFDRTLARGPQTTTWIWNLHALAHDFDT 60
Synechocystis 6803 MTISPPEREAKAKVSVDNNPVPTSFEKWGKPGHFDRTLARGPKTTTWIWNLHANAHDFDS 60
                   ********* *.:. **::*********.**********:**:********** *****:

S. leopoliensis    HTSDLEDISRKIFSAHFGHLAVIFIWLSGAYFHGARFSNFSGWLADPTHVKPSAQVVWPI 120
T. vestitus        HTSDLEDISRKIFSAHFGHLAVVFIWLSGMYFHGAKFSNYEAWLADPTGIKPSAQVVWPI 120
S. lividus         HTSDLEDISRKIFSAHFGHLAVVFIWLSGMYFHGAKFSNYEAWLADPTGIKPSAQVVWPI 120
Synechocystis 6803 QTSDLEDVSRKIFSAHFGHLAVVFVWLSGMYFHGAKFSNYEGWLADPTHIKPSAQVVWPI 120
                   :******:**************:*:**** *****:***:..****** :**********

S. leopoliensis    FGQEILNGDVGGGFHGIQITSGLFQLWRASGYTNEFQLYVTAIGALVMAGLMLFAGWFHY 180
T. vestitus        VGQGILNGDVGGGFHGIQITSGLFQLWRASGITNEFQLYCTAIGGLVMAGLMLFAGWFHY 180
S. lividus         VGQGILNGDVGGGFHGIQITSGLFQLWRASGITNEFQLYCTAIGGLVMAGLMLFAGWFHY 180
Synechocystis 6803 VGQGILNGDVGGGFHGIQITSGLFYLWRASGFTDSYQLYCTAIGGLVMAALMLFAGWFHY 180
                   .** ******************** ****** *:.:*** ****.****.**********

S. leopoliensis    HKAAPKLEWFQNVESMLNHHLAGLLGLGSLSWAGHQIHVSLPVNKLLDAIDAGEPLVLNG 240
T. vestitus        HKRAPKLEWFQNVESMLNHHLAGLLGLGSLAWAGHQIHVSLPINKLLDAGV--------- 231
S. lividus         HKRAPKLEWFQNVESMLNHHLAGLLGLGSLGWAGHQIHVSLPINQLLDAGV--------- 231
Synechocystis 6803 HVKAPKLEWFQNVESMMNHHLAGLLGLGSLGWAGHQIHVSMPINKLLDAGV--------- 231
                   *  *************:*************.*********:*:*:****

S. leopoliensis    KTIASAADIPLPHEFL-DVSLISQLFPG----FEAGVKAFFTLNWSAYADFLTFKGGLNP 295
T. vestitus        ----AAKDIPLPHEFILNPSLMAELYPKVDWGFFSGVIPFFTFNWAAYSDFLTFNGGLNP 287
S. lividus         ----AAKDIPLPHEFILNPSLMAELYPNINWGVFSGVIPFFTFNWAAYSDFLTFKGGLNP 287
Synechocystis 6803 ----APKDIPLPHEFILEPSKMAELYPSF----AQGLTPFFTLNWGVYSDFLTFKGGLNP 283
                       :  ********: : * :::*:*        *:  ***:**..*:*****:*****

S. leopoliensis    VTGGLWLTDTAHHHLAIAVLFIVAGHMYRTNWGIGHSLKEILEAHKGPFTGQGHKGLYEI 355
T. vestitus        VTGGLWLSDTAHHHLAIAVLFIIAGHMYRTNWGIGHSLKEILEAHKGPFTGAGHKGLYEV 347
S. lividus         VTGGLWLSDTAHHHVAIAVLFIIAGHMYRTNWGIGHSLKEILEAHKGPFTGTGHKGLYEV 347
Synechocystis 6803 VTGGLWLSDTAHHHLAIAVLFIIAGHMYRTNWGIGHSMKEILEAHKGPFTGEGHKGLYEI 343
                   *******:******:*******:**************:************* *******:

S. leopoliensis    LTTSWHAQLSINLAILGSISIIVAHHMYAMPPYPYLATDYPTMLSLFTHHIWIGGFLIVG 415
T. vestitus        LTTSWHAQLAINLAMMGSLSIIVAQHMYAMPPYPYLATDYPTQLSLFTHHMWIGGFLVVG 407
S. lividus         LTTSWHAQLAINLAMMGSLSIIVAQHMYAMPPYPYLATDYPTQLSLFTHHMWIGGFLIVG 407
Synechocystis 6803 LTTSWHAQLAINLALLGSLTIIVAQHMYAMPPYPYQAIDYATQLSLFTHHMWIGGFLIVG 403
                   *********:****::**::****:********** * ** * *******:******:**

S. leopoliensis    AGAHAAIFMVRDYDPAKNVDNLLDRVLRHRDAIISHLNWVCIWLGFHSFGLYIHNDTMRA 475
T. vestitus        GAAHGAIFMVRDYDPAMNQNNVLDRVLRHRDAIISHLNWVCIFLGFHSFGLYVHNDTMRA 467
S. lividus         GAAHGAIFFVRDYDPAVNQNNVLDRVLRHRDAIISHLNWVCIFLGFHSFGLYVHNDTMRA 467
Synechocystis 6803 AGAHGAIFMVRDYDPAKNVNNLLDRMLRHRDAIISHLNWVCIFLGFHSFGLYIHNDTMRA 463
                   ..**.***:******* * :*:***:****************:*********:*******

S. leopoliensis    LGRPQDMFSDSAIQLQPIFAQWIQNIHALAPGNTAPNALASVSQVFGGDVVAVGGKVAAA 535
T. vestitus        FGRPQDMFSDTGIQLQPVFAQWVQNLHTLAPGGTAPNAAATASVAFGGDVVAVGGKVAMM 527
S. lividus         FGRPQDMFSDTGIQLQPVFAQWVQHLHTLAPGGTAPNAAATASVAFGGDVVAVGGKVAMM 527
Synechocystis 6803 LGRPQDMFSDTAIQLQPIFAQWVQHLHTLAPGATAPNALATASYAFGGETIAVAGKVAMM 523
                   :*********:.*****:****:*::*:**** ***** *:.* .***:.:**.****

S. leopoliensis    PIVLGTADFMVHHIHAFTIHVTALILLKGVLYARSSRLVPDKANLGFRFPCDGPGRGGTC 595
T. vestitus        PIVLGTADFMVHHIHAFTIHVTVLILLKGVLFARSSRLIPDKANLGFRFPCDGPGRGGTC 587
S. lividus         PIALGTADFLVHHIHAFTIHVTVLILLKGVLFARSSRLIPDKANLGFRFPCDGPGRGGTC 587
Synechocystis 6803 PITLGTADFMVHHIHAFTIHVTALILLKGVLYARSSRLVPDKANLGFRFPCDGPGRGGTC 583
                   **.******:************.********:******:*********************

S. leopoliensis    QVSGWDHVFLGLFWMYNSLSIVIFHYSWKMQSDVWGSVLPDGSVAHIANGNFAQSALTIN 655
T. vestitus        QVSGWDHVFLGLFWMYNCISVVIFHFSWKMQSDVWGTVAPDGTVSHITGGNFAQSAITIN 647
S. lividus         QVSGWDHVFLGLFWMYNCISVVIFHFSWKMQSDVWGTVAPDGTVSHITGGNFAQSAITIN 647
Synechocystis 6803 QVSGWDHVFLGLFWMYNSLSIVIFHFSWKMQSDVWGTVSPDGSVTHVTLGNFAQSAITIN 643
                   *****************.:*:****:**********:* ***:*:*:: *******:***

S. leopoliensis    GWLRDFLWAQASQVITSYGSSTSAYGLLFLGAHFVWAFSLMFLFSGRGYWQELIESIVWA 715
T. vestitus        GWLRDFLWAQASQVIGSYGSALSAYGLLFLGAHFIWAFSLMFLFSGRGYWQELIESIVWA 707
S. lividus         GWLRDFLWAQASQVIGSYGSALSAYGLLFLGAHFVWAFSLMFLFSGRGYWQELIESIVWA 707
Synechocystis 6803 GWLRDFLWAQAANVINSYGSALSAYGIMFLAGHFVFAFSLMFLFSGRGYWQELIESIVWA 703
                   ***********::** ****: ****::**..**::************************

S. leopoliensis    HNKLKVAPAIQPRALSIIQGRAVGVAHYLLGGIVTTWSFFLARIIAVG 763
T. vestitus        HNKLKVAPAIQPRALSIIQGRAVGVAHYLLGGIATTWAFFLARIISVG 755
S. lividus         HNKLKVAPAIQPRALSIIQGRAVGVAHYLLGGIATTWAFFLARIISVG 755
Synechocystis 6803 HNKLNVAPAIQPRALSIIQGRAVGVAHYLLGGIVTTWAFFLARSLSIG 751
                   ****:****************************.***:***** :::*
```

### PsaB

```
S. leopoliensis    MATKFPKFSQDLAQDPTTRRIWYGIATAHDFESHDGMTEENLYQKIFASHFGHLAIIFLW 60
T. vestitus        MATKFPKFSQDLAQDPTTRRIWYAIAMAHDFESHDGMTEENLYQKIFASHFGHLAIIFLW 60
S. lividus         MATKFPKFSQDLAQDPTTRRIWYAIATAHDFESHDGMTEENLYQKIFASHFGHLAIIFLW 60
Synechocystis 6803 MATKFPKFSQDLAQDPTTRRIWYGIATAHDFETHDGMTEENLYQKIFASHFGHIAIIFLW 60
                   ***********************.** *****:********************:******

S. leopoliensis    VSGNLFHVAWQGNFEQWSQDPLHVRPIAHAIWDPHFGQGAIDAFTQAGASSPVNVAYSGV 120
T. vestitus        VSGSLFHVAWQGNFEQWVQDPVNTRPIAHAIWDPQFGKAAVDAFTQAGASNPVDIAYSGV 120
S. lividus         VSGSLFHVAWQGNFEQWIQDPLNTRPIAHAIWDPQFGKAAVDAFTQAGASSPVDIAYSGV 120
Synechocystis 6803 TSGTLFHVAWQGNFEQWIKDPLNIRPIAHAIWDPHFGEGAVNAFTQAGASNPVNIAYSGV 120
                   .**.************* :**:: **********:**:.*::********.**::*****

S. leopoliensis    YHWWYTIGMRTNGDLYQGSIFLLILSALFLFAGWLHLQPKFRPSLSWFKNAESRLNHHLA 180
T. vestitus        YHWWYTIGMRTNGDLYQGAIFLLILASLALFAGWLHLQPKFRPSLSWFKNAESRLNHHLA 180
S. lividus         YHWWYTIGMRTNGDLYQGAIFLLVLASLALFAGWLHLQPKFRPSLSWFKNAESRLNHHLA 180
Synechocystis 6803 YHWFYTIGMTTNQELYSGAVFLLVLASLFLFAGWLHLQPKFRPSLAWFKNAESRLNHHLA 180
                   ***:***** ** :**.*::***:*::* ****************:**************

S. leopoliensis    GLFGFSSLAWTGHLVHVAIPEARGQHVGWDNFLSTLPHPAGLAPFFTGNWSVYAENPDTA 240
T. vestitus        GLFGVSSLAWAGHLIHVAIPESRGQHVGWDNFLSTMPHPAGLAPFFTGNWGVYAQNPDTA 240
S. lividus         GLFGVSSLAWAGHLIHVAIPESRGQHVGWDNFLSTMPHPAGLAPFFTGNWGVYAQNPDTA 240
Synechocystis 6803 GLFGVSSLAWAGHLVHVAIPEARGQHVGWDNFLSTPPHPAGLMPFFTGNWGVYAADPDTA 240
                   ****.*****:***:******:************* ****** *******.*** :****

S. leopoliensis    SHAFGTAEGAGTAILTFLGGFHPQTEALWLTDIAHHHLAIAVIFIIAGHMYRTNFGIGHS 300
T. vestitus        SHVFGTAQGAGTAILTFLGGFHPQTESLWLTDMAHHHLAIAVLFIVAGHMYRTQFGIGHS 300
S. lividus         SHVFGTSQGAGSAILTFLGGFHPQTESLWLTDMAHHHLAIAVLFIVAGHMYRTQFGIGHS 300
Synechocystis 6803 GHIFGTSEGAGTAILTFLGGFHPQTESLWLTDIAHHHLAIAVIFIIAGHMYRTNWGIGHS 300
                   .* ***::***:**************:*****:*********:**:*******::*****

S. leopoliensis    IKEILEAHKPP----AGGLGAGHKGLYETLNNSLHFQLALALASLGVVTSLVAQHMYSMP 356
T. vestitus        IKEMMDAKDFFGTKVEGPFNMPHQGIYETYNNSLHFQLGWHLACLGVITSLVAQHMYSLP 360
S. lividus         IKEMMDAKDFFGTKVEGPFNMPHQGIYETYNNSLHFQLGWHLACLGVITSLVAQHMYSLP 360
Synechocystis 6803 IKEILNAHKG------PLTGAGHTNLYDTINNSLHFQLGLALASLGVITSLVAQHMYSLP 354
                   ***:::*:.          .  * .:*:* ********.  **.***:**********:*

S. leopoliensis    PYAFIAKDYTTMAALYTHHQYIATFIMCGAFAHGAIFLIRDYDPEANKNNVLARVLEHKE 416
T. vestitus        PYAFIAQDHTTMAALYTHHQYIAGFLMVGAFAHGAIFLVRDYDPAQNKGNVLDRVLQHKE 420
S. lividus         PYAFIAQDHTTMAALYTHHQYIAGFLMVGAFAHGAIFLVRDYDPVQNKGNVLDRVLQHKE 420
Synechocystis 6803 SYAFIAQDHTTQAALYTHHQYIAGFLMVGAFAHGAIFFVRDYDPVANKDNVLARMLEHKE 414
                    *****:*:** *********** *:* *********::*****  **.*** *:*:***

S. leopoliensis    AIISHLSWVSLFLGFHTLGLYVHNDVVVAFGTPEKQILIEPVFAQFVQAASGKALYGFNV 476
T. vestitus        AIISHLSWVSLFLGFHTLGLYVHNDVVVAFGTPEKQILIEPVFAQFIQAAHGKLLYGFDT 480
S. lividus         AIISHLSWVSLFLGFHTLGLYVHNDVVVAFGTPEKQILIEPVFAQFIQAAHGKLLYGFDT 480
Synechocystis 6803 ALISHLSWVSLFLGFHTLGLYVHNDVVVAFGTPEKQILIEPVFAQWIQATSGKALYGFDV 474
                   *:*******************************************::**: ** ****:.

S. leopoliensis    LLANADSAATAA---SLGTYLPNWLDAINSGKTALFLPIGPGDFLVHHAIALGLHTTTLI 533
T. vestitus        LLSNPDSIASTAWPNYGNVWLPGWLDAINSGTNSLFLTIGPGDFLVHHAIALGLHTTTLI 540
S. lividus         LLSNPDSIAATAWPNYGNVWLPGWLDAINSGNNSLFLTIGPGDFLVHHAIALGLHTTTLI 540
Synechocystis 6803 LLSNPDSIAST----TGAAWLPGWLDAINSGTNSLFLTIGPGDFLVHHAIALGLHTTALI 530
                   **:* ** *::       .:**.********..:*** *******************:**

S. leopoliensis    LVKGALDARGSKLMPDKKDFGYSFPCDGPGRGGTCDISAWDAFYLAVFWALNTVGWVTFY 593
T. vestitus        LVKGALDARGSKLMPDKKDFGYAFPCDGPGRGGTCDISAWDAFYLAMFWMLNTIGWVTFY 600
S. lividus         LVKGALDARGSKLMPDKKDFGYAFPCDGPGRGGTCDISAWDAFYLAMFWMLNTIGWVTFY 600
Synechocystis 6803 LIKGALDARGSKLMPDKKDFGYSFPCDGPGRGGTCDISAWDAFYLAMFWMLNTLGWLTFY 590
                   *:********************:***********************:** ***:**:***

S. leopoliensis    WHWKNLTVWQGNVAQFNESSTYLMGWLRDYLWLNSSQLINGYNPFGTNNLSVWSWMFLFG 653
T. vestitus        WHWKHLGVWEGNVAQFNESSTYLMGWLRDYLWLNSSQLINGYNPFGTNNLSVWAWMFLFG 660
S. lividus         WHWKHLGVWEGNVAQFNENSTYLMGWLRDYLWLNSSQLINGYNPFGTNNLSVWAWMFLFG 660
Synechocystis 6803 WHWKHLGVWSGNVAQFNENSTYLMGWFRDYLWANSAQLINGYNPYGVNNLSVWAWMFLFG 650
                   ****:* **.********.*******:***** **:********:*.******:******

S. leopoliensis    HLIWATGFMFLISWRGYWQELIETIVWAHQRTPLANIVGWKDKPVALSIVQARVVGLAHF 713
T. vestitus        HLVWATGFMFLISWRGYWQELIETLVWAHERTPLANLVRWKDKPVALSIVQARLVGLAHF 720
S. lividus         HLVWATGFMFLISWRGYWQELIETLVWAHERTPLANLVRWKDKPVALSIVQARLVGLAHF 720
Synechocystis 6803 HLVWATGFMFLISWRGYWQELIETIVWAHERTPLANLVRWKDKPVALSIVQARLVGLAHF 710
                   **:*********************:****:******:* **************:******

S. leopoliensis    TVGYFLTYAAFLIASTAGKFG 734
T. vestitus        SVGYILTYAAFLIASTAAKFG 741
S. lividus         SVGYVLTYAAFLIASTASKYG 741
Synechocystis 6803 TVGYVLTYAAFLIASTAGKFG 731
                   :***.************.*:*
```

### PsaC

```
S. leopoliensis    MSHSVKIYDTCIGCTQCVRACPLDVLEMVPWDGCKAGQIAASPRTEDCVGCKRCETACPT       60
T. vestitus        MAHTVKIYDTCIGCTQCVRACPTDVLEMVPWDGCKAGQIASSPRTEDCVGCKRCETACPT       60
S. lividus         MAHTVKIYDTCIGCTQCVRACPTDVLEMVPWDGCKAGQIASSPRTEDCVGCKRCETACPT       60
Synechocystis 6803 MSHSVKIYDTCIGCTQCVRACPLDVLEMVPWDGCKAAQIASSPRTEDCVGCKRCETACPT       60
                   *:*:****************** *************.***:*******************

S. leopoliensis    DFLSIRVYLGAETTRSMGLAY   81
T. vestitus        DFLSIRVYLGAETTRSMGLAY   81
S. lividus         DFLSIRVYLGAETTRSMGLAY   81
Synechocystis 6803 DFLSIRVYLGAETTRSMGLAY   81
                   *********************
```

### PsaD

```
S. leopoliensis    MAETLTGKTPVFGGSTGGLLKSAETEEKYAITWTSTKEQVFELPTGGAAVMHEGDNLLYF 60
T. vestitus        -MTTLTGQPPLYGGSTGGLLSAADTEEKYAITWTSPKEQVFEMPTAGAAVMREGENLVYF 59
S. lividus         -MTTLTGQPPLYGGSTGGLLSKADIEEKYAITWTSPKEQVFEMPTAGAAVMREGENLVYL 59
Synechocystis 6803 -MTELSGQPPKFGGSTGGLLSKANREEKYAITWTSASEQVFEMPTGGAAIMNEGENLLYL 59
                       *:*: * :********. *: ********** .*****:**.***:*.**:**:*:

S. leopoliensis    ARKEQALALGTQLRTKFKPKIESYKIYRVFPGGDVEYLHPKDGVFPEKVNEGRSFAGKVD 120
T. vestitus        ARKEQCLALAAQQL--RPRKINDYKIYRIFPDGETVLIHPKDGVFPEKVNKGREAVNSVP 117
S. lividus         ARKEQCLALAAQQL--RPRKINDYKIYRIFPDGETVLLHPKDGVFPEKVNQGREAVNTVP 117
Synechocystis 6803 ARKEQCLALGTQLRTKFKPKIQDYKIYRVYPSGEVQYLHPADGVFPEKVNEGREAQGTKT 119
                   *****.***.:*       **:.*****::*.*:.  :** *********:**.  ..

S. leopoliensis    RRIGQNPNPATIKFTGKQPYTA 142
T. vestitus        RSIGQNPNPSQLKFTGKKPYDP 139
S. lividus         RSIGQNPNPAQLKFSGKKPYDP 139
Synechocystis 6803 RRIGQNPEPVTIKFSGKAPYEV 141
                   * *****:*  :**:** **
```

### PsaE

```
S. leopoliensis    MAIARGDKVRILRPESYWFNEVGTVASVDQ-SGIKYPVVVRFEKVNYNGFSGSDGGVNTN 59
T. vestitus        -MVQRGSKVKILRPESYWYNEVGTVASVDQTPGVKYPVIVRFDKVNYTGYSGSASGVNTN 59
S. lividus         -MVQRGSKVRILRPESYWYNEIGTVASVDQ-SGIKYPVIVRFDKVNYTGFSGSAGGVNTN 58
Synechocystis 6803 MALNRGDKVRIKRTESYWYGDVGTVASVEK-SGILYPVIVRFDRVNYNGFSGSASGVNTN 59
                     : **.**:* * ****:.::******::  *: ***:***::***.*:*** .*****

S. leopoliensis    NFAEAELQVVAAAAKK- 75
T. vestitus        NFALHEVQEVAPPKKGK 76
S. lividus         NFALSEVEEVSAPKKGK 75
Synechocystis 6803 NFAENELELVQAAAK-- 74
                   ***  *:: *    *
```

### PsaF

```
S. leopoliensis    MRRLFAVVLAACLWLGFAPQASA-DVAGLTPCSESPRFIQRAEAAAT-----PQAKARFE 54
T. vestitus        MRRFLALLLVLTLWLGFTPLASA-DVAGLVPCKDSPAFQKRAAAAVNTTADPASGQKRFE 59
S. lividus         MRRLLALLLVLSLWIGFTPLASA-DVAGLVPCKDSPAFQKRAAAAVNTTADPASGQKRFE 59
Synechocystis 6803 MKHLLALLLAFTLWFNFAPSASADDFANLTPCSENPAYLAKSKNFLNTTNDPNSGKIRAE 60
                   *::::*::*.  **:.*:* *** *.*.*.**.:.* :  ::    .      ..: * *

S. leopoliensis    NYSQALCGADGLPHLIVDGRLDHAGDFIIPSLLFLYIAGWIGWVGRSYLQAIKSDKDAAG 114
T. vestitus        RYSQALCGEDGLPHLVVDGRLSRAGDFLIPSVLFLYIAGWIGWVGRAYLIAVRNSGEANE 119
S. lividus         RYSQALCGEDGLPHLVVDGRLSRAGDFLIPSALFLFITGWIGWVGRAYLIAVRNSGEANE 119
Synechocystis 6803 RYASALCGPEGYPHLIVDGRFTHAGDFLIPSILFLYIAGWIGWVGRSYLIEIRESKNPEM 120
                   .*:.**** :* ***:****: :****:*** ***:*:********:**  ::.. :

S. leopoliensis    KEIVIDVPLAVKFSLTGFAWPLAAFQEFSSGKLLAKADEITVSPR 159
T. vestitus        KEIIIDVPLAIKCMLTGFAWPLAALKELASGELTAKDNEITVSPR 164
S. lividus         KEIIIDVPLAIKCMLTGFAWPLAALKELASGELTAKDNEITVSPR 164
Synechocystis 6803 QEVVINVPLAIKKMLGGFLWPLAAVGEYTSGKLVMKDSEIPTSPR 165
                   :*::*:****:*  * ** *****. * :**:*  * .** .***
```

### PsaI

```
S. leopoliensis    MSGDFAAAFLPTIFVPLVGLGLPAVLMSLLFTYIESEA-- 38
T. vestitus        MMGSYAASFLPWIFIPVVCWLMPTVVMGLLFLYIEGEA-- 38
S. lividus         MMGSYAASFLPWIFIPVVCWLMPVVVMGLLFLYIEGDAQA 40
Synechocystis 6803 MDGSYAASYLPWILIPMVGWLFPAVTMGLLFIHIESEGEG 40
                   * *.:**::** *::*:*   :*.* *.*** :**.:.
```

### PsaJ

```
S. leopoliensis    MLAMDGLKRYLSSAPILATIWFAITAGILIEFNRFFPDLLFHPL 44
T. vestitus        ---MKHFLTYLSTAPVLAAIWMTITAGILIEFNRFYPDLLFHPL 41
S. lividus         -MQTKHLLTYLSTAPVLAALWMTITAGILIEFNRFYPDLLFHPL 43
Synechocystis 6803 ---MDGLKSFLSTAPVMIMALLTFTAGILIEFNRFYPDLLFHP- 40
                       . :  :**:**::    :::***********:*******
```

### PsaK

```
S. leopoliensis (PsaK1) ----------MNPTTVEWNANVAAIMITANLFAIAIGYFAIRNRGVGPALPVPLPAIFSG 50
S. leopoliensis (PsaK2) ----MLPVLAAIPQTVAWSPKVALVMILSNIVAIAIGKATIKIQNAGPALPS--PQLFGG 54
T. vestitus             -------MVLATLPDTTWTPSVGLVVILCNLFAIALGRYAIQSRGKGPGLPIALPALFEG 53
S. lividus              -------MVLATLPDTTWSPAVGLVVILCNLFAIAIGRYAIQSRGKGPGLPVSLPALFEG 53
Synechocystis 6803      MFNTALLLAQASPTTAGWSLSVGIIMCLCNVFAFVIGYFAIQKTGKGKDLALPQLASKKT 60
                                       . *.  *. ::  .*:.*:.:*  :*:  . *  *

S. leopoliensis (PsaK1) FGLPELLATASFGHLLGAGFVLGLAQAGLL 80
S. leopoliensis (PsaK2) FGLPAVLATASFGHILGIGVILGLANIGNL 84
T. vestitus             FGLPELLATTSFGHLLAAGVVSGLQYAGAL 83
S. lividus              FGLPELLATTSFGHLLAAGVVSGLQYSGAL 83
Synechocystis 6803      FGLPELLATMSFGHILGAGMVLGLASSGIL 90
                        **** :*** ****:*. *.: **   * *
```

### PsaL

```
S. leopoliensis    ---MAQDVIANGGTPEIGNLATPINSSPFTRTFINALPIYRRGLSSNRRGLEIGMAHGFL 57
T. vestitus        --MAEELVKPYNGDPFVGHLSTPISDSGLVKTFIGNLPAYRQGLSPILRGLEVGMAHGYF 58
S. lividus         --MADELVKPYNGDPFAGHLSTPISDSGLVKTFISNLPAYRQGLSPILRGLEVGMAHGYF 58
Synechocystis 6803 MAESNQVVQAYNGDPFVGHLSTPISDSAFTRTFIGNLPAYRKGLSPILRGLEVGMAHGYF 60
                        : *   .* *  *:*:***..* :.:***. ** **:***   ****:*****::

S. leopoliensis    LYGPFSILGPLRNTETAGSAGLLATVGLVVILTVCLSLYGNAGSGPSAAESTVTTPNPPQ 117
T. vestitus        LIGPWVKLGPLRDSDVANLGGLISGIALILVATACLAAYGLVSFQKG--------GSSSD 110
S. lividus         LIGPWVKLGPLRDSDVANLGGLISGITLILLATACLAAYGLVSFQKA--------SSSGD 110
Synechocystis 6803 LIGPWTLLGPLRDSEYQYIGGLIGALALILVATAALSSYGLVTFQGE--------QGSGD 112
                   * **:  *****:::    .**:. : *::: *..*: ** .              .  :

S. leopoliensis    ELFTKEGWSEFTSGFILGGLGGAFFAFYLASTPYVQPLVKIAAGVWSVH 166
T. vestitus        PLKTSEGWSQFTAGFFVGAMGSAFVAFFLLENFSVV--DGIMTGLFN-- 155
S. lividus         ALKTGEGWSQFTAGFFVGAMGGAFVAFFLLENFAVV--DGIMKGLFN-- 155
Synechocystis 6803 TLQTADGWSQFAAGFFVGGMGGAFVAYFLLENLSVV--DGIFRGLFN-- 157
                    * * :***:*::**::*.:*.**.*::* ..  *     *  *::.
```

### PsaM

```
S. leopoliensis    --MTDTQVFVALLLALVPAVLAYRLGTELYR 29
Synechocystis 6803 MALSDTQILAALVVALLPAFLAFRLSTELYK 31
T. vestitus        MALTDTQVYVALVIALLPAVLAFRLSTELYK 31
S. lividus         MALTDTQVYIALVIALLPAVLAFRLSAELYK 31
                     ::***:  **::**:**.**:**.:***:
```

### PsaX

```
T. vestitus MSTMATKSAKPTYAFRTFWAVLLLAINFLVAAYYFGILK 39
S. lividus  ---MATKSAKPTYTFRTFWAVLLLAINFLVAAYYFGILK 36
               **********:*************************
```

**Fig. S5.** Multiple sequence alignments of PSI subunits from *S. leopoliensis*, *T. vestitus*, and *Synechocystis* 6803. Note that of these organisms, PsaX is found only in *T. vestitus* and *S. lividus*. Sequence identities were calculated using Clustal Omega (6, 7).

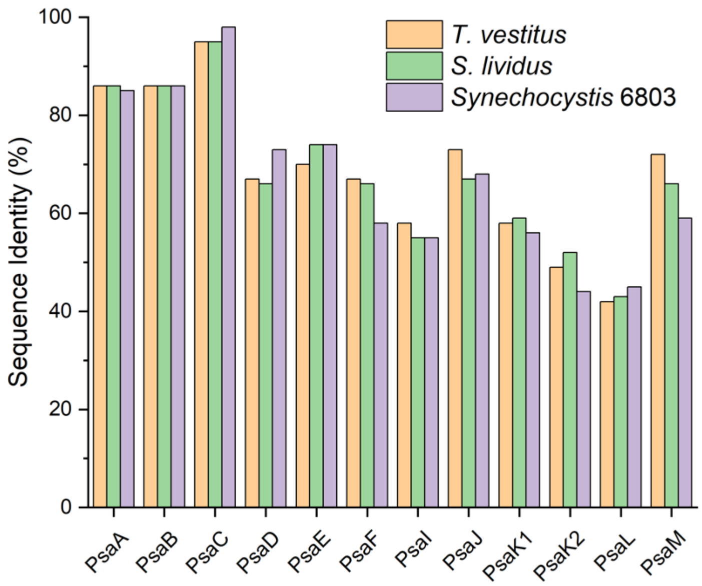


**Fig. S6.** Sequence identities of PSI subunits in *S. leopoliensis* compared to homologous subunits in *T. vestitus*, *S. lividus*, and *Synechocystis* 6803. Sequence identities were calculated using Clustal Omega.

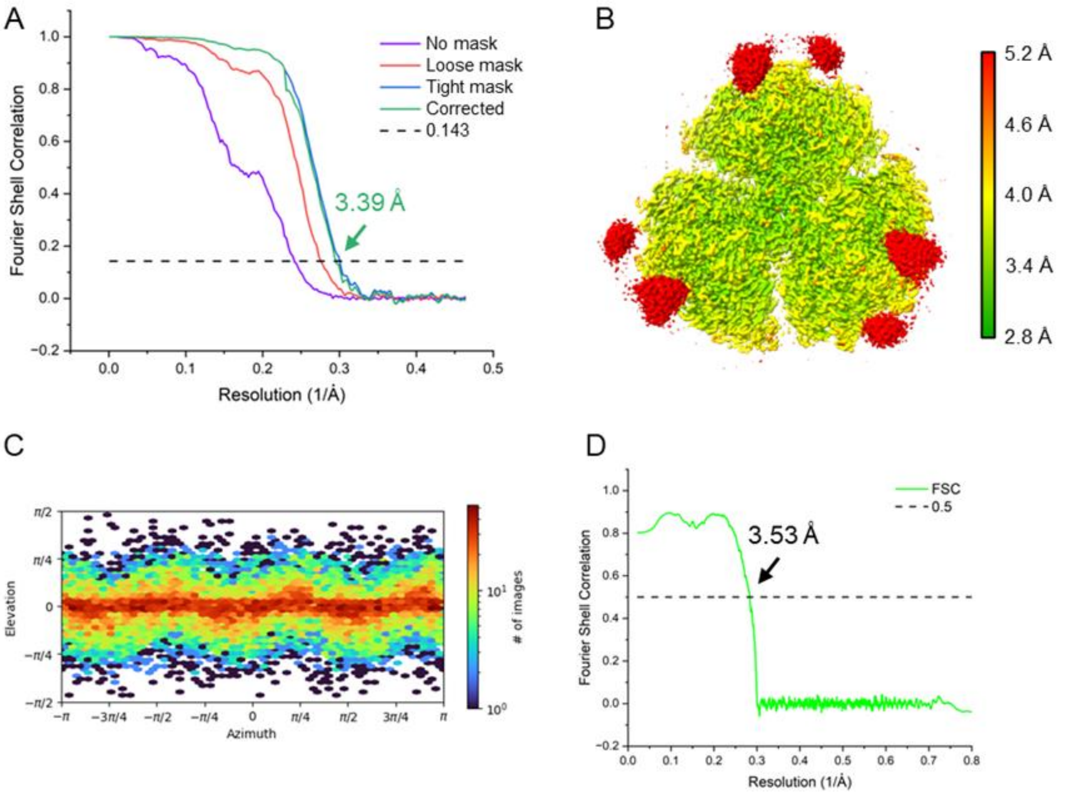


**Fig. S7.** Pt-bPSI$^{tri}_{T.v.}$ cryo-EM resolution and data completeness. (*A*) Map-to-map Fourier Shell Correlation determining the reported global resolution. (*B*) local resolution map. (*C*) Angular distribution of cryo-EM data. (*D*) Map-to-model Fourier Shell Correlation.

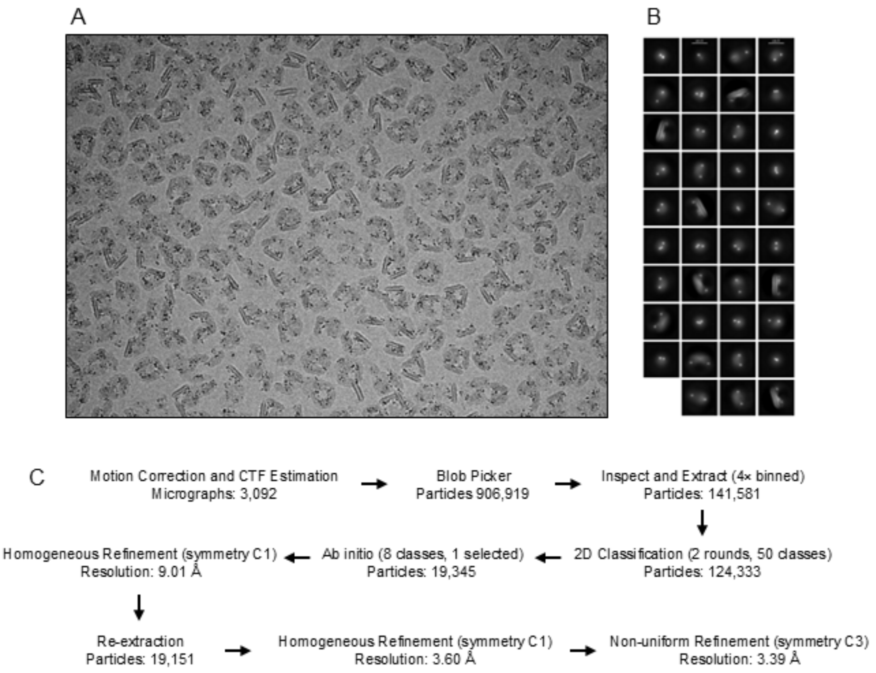


**Fig. S8.** Pt-bPSI$^{tri}_{T.v.}$ cryo-EM data quality and workflow. (*A*) Example micrograph. (*B*) 2D classes of accepted particles generated in CryoSPARC. (*C*) The workflow for cryo-EM data processing in CryoSPARC.

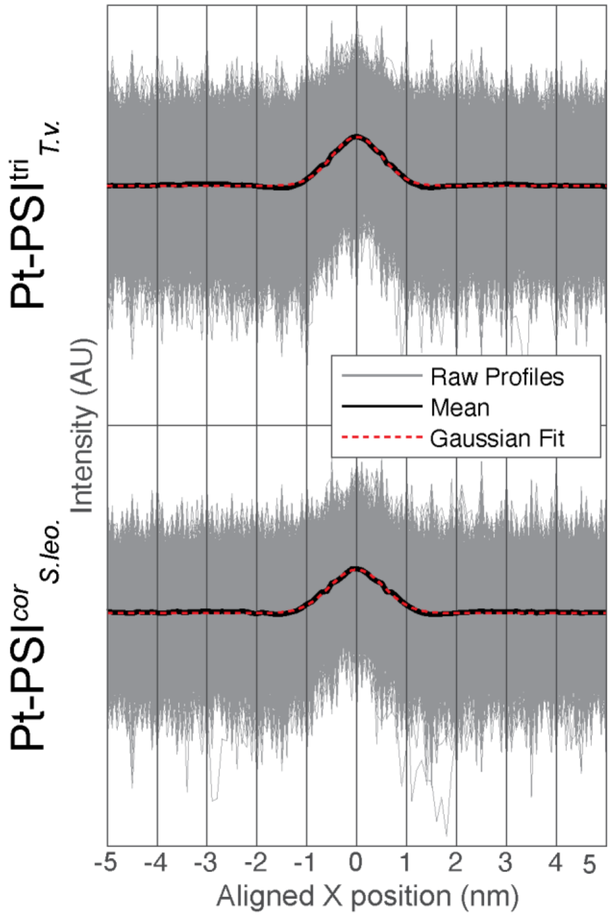


**Fig. S9.** PtNP profiles and mean traces. Overlaid traces from computationally segmented/located PtNPs from Pt-PSI$^{tri}_{T.v.}$ (top) and Pt-PSI$^{cor}_{S.leo.}$ (bottom) micrographs. Data sets comprise 7,080 and 5,188 particle traces for Pt-PSI$^{tri}_{T.v.}$ and Pt-PSI$^{cor}_{S.leo}$ respectively, from 10 micrographs each. See **Materials and Methods** for a description of computational pipeline and code availability.

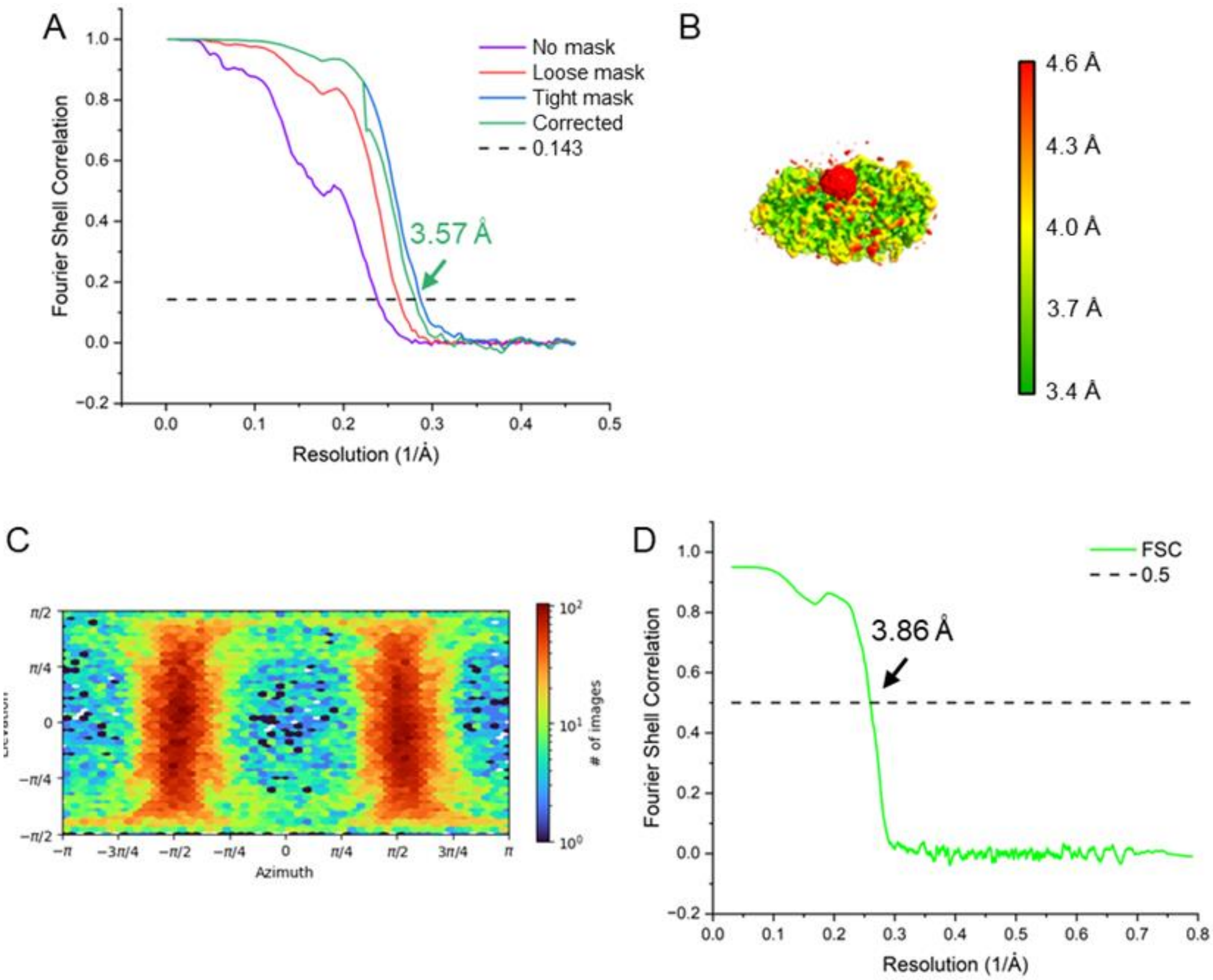


**Fig. S10.** Pt-bPSI$^{cor}_{S.leo.}$ cryo-EM resolution and data completeness. (*A*) Map-to-map Fourier Shell Correlation determining the reported global resolution. (*B*) local resolution map. (*C*) Angular distribution of cryo-EM data. (*D*) Map-to-model Fourier Shell Correlation.

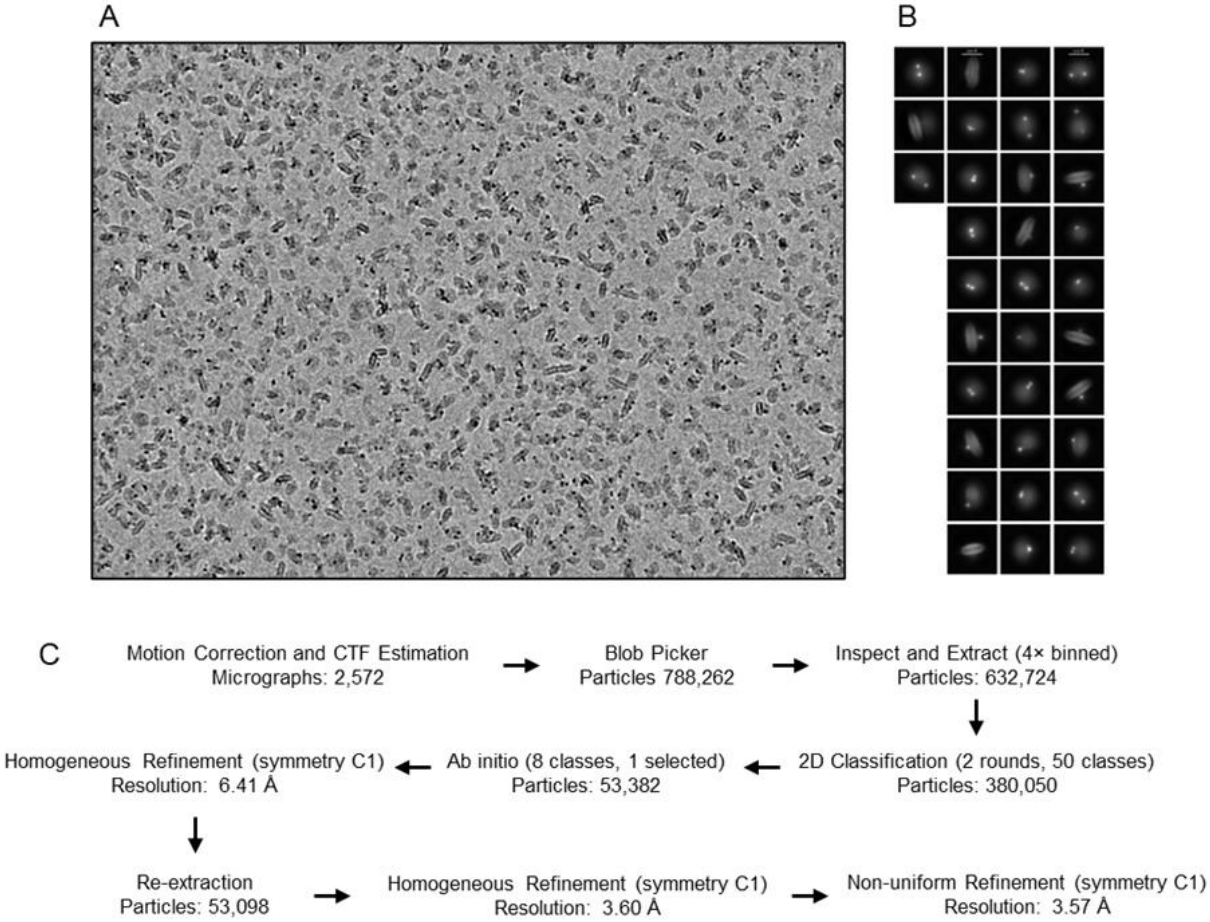


**Fig. S11.** Pt-bPSI$^{cor}_{\textit{S.leo.}}$ cryo-EM data quality and workflow. (*A*) Example micrograph. (*B*) 2D classes of accepted particles generated in CryoSPARC. (*C*) The workflow for cryo-EM data processing in CryoSPARC.

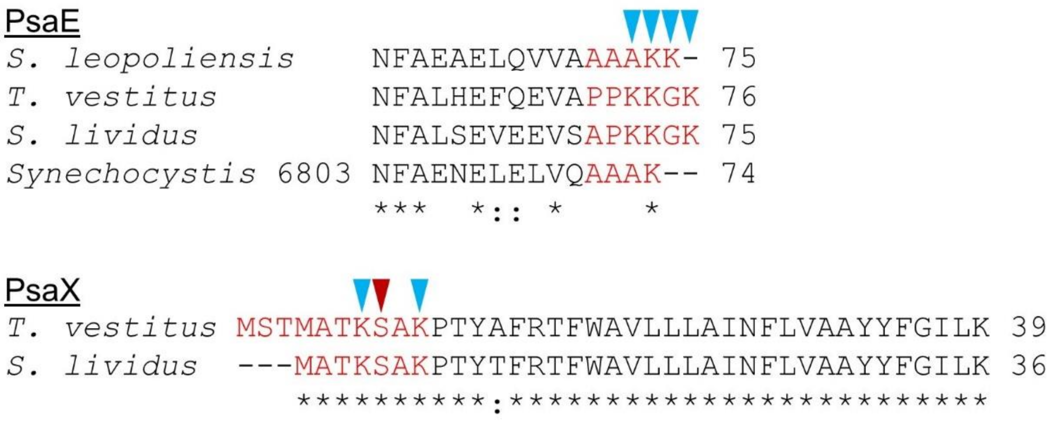


**Fig. S12.** Unmodelled residues of PsaE and PsaX in multiple cyanobacterial strains. Abridged sequence alignments of PsaE (top) and PsaX (bottom) showing residues which are not modelled in the Pt-bPSI$^{tri}_{T.v.}$ structure due to a lack of signal in the cryo-EM map in red text. Blue arrows indicate positions in these regions which can contain positive residues, while the red arrow indicates the position with a polar residue—those which may interact with PtNPs.

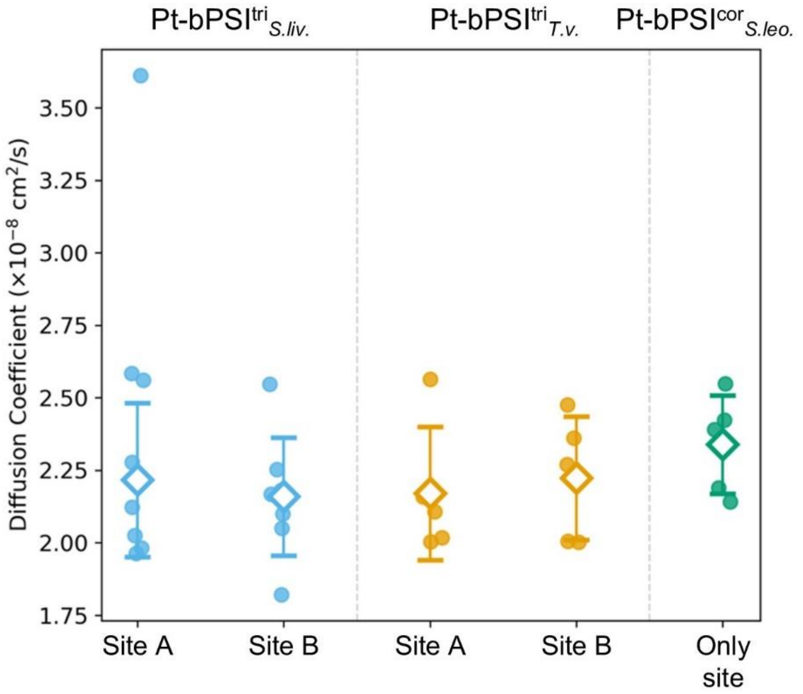


**Fig. S13.** PtNP diffusion coefficients on PSI surfaces. Diffusion coefficients of PtNPs on PSI surfaces were extracted from mean squared displacement slopes fit over the first 1 µs of simulation time. Points represent independent replicas, diamonds indicate the means, and error bars denote ±1 standard deviation.

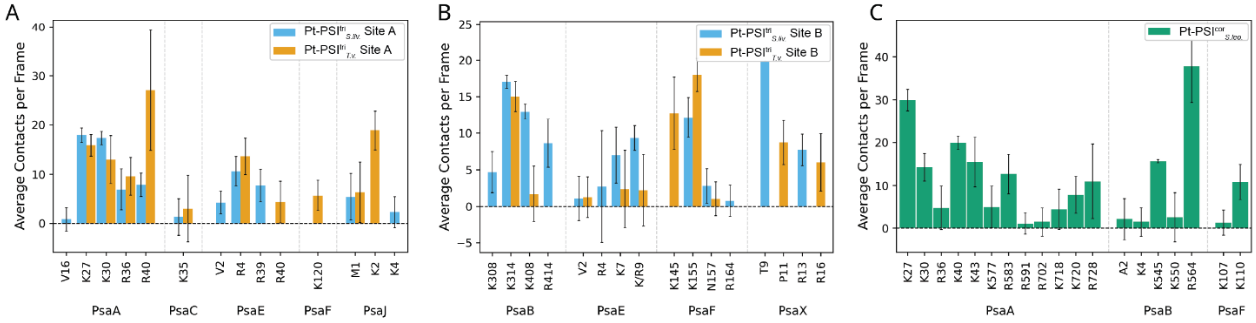


**Fig. S14.** PSI residues contributing to PtNP stabilization. Bar plots show the average PtNP–residue contact frequency per frame for all residues identified from the contact analysis. These residues include both dominant anchoring residues and lower-frequency interaction sites.

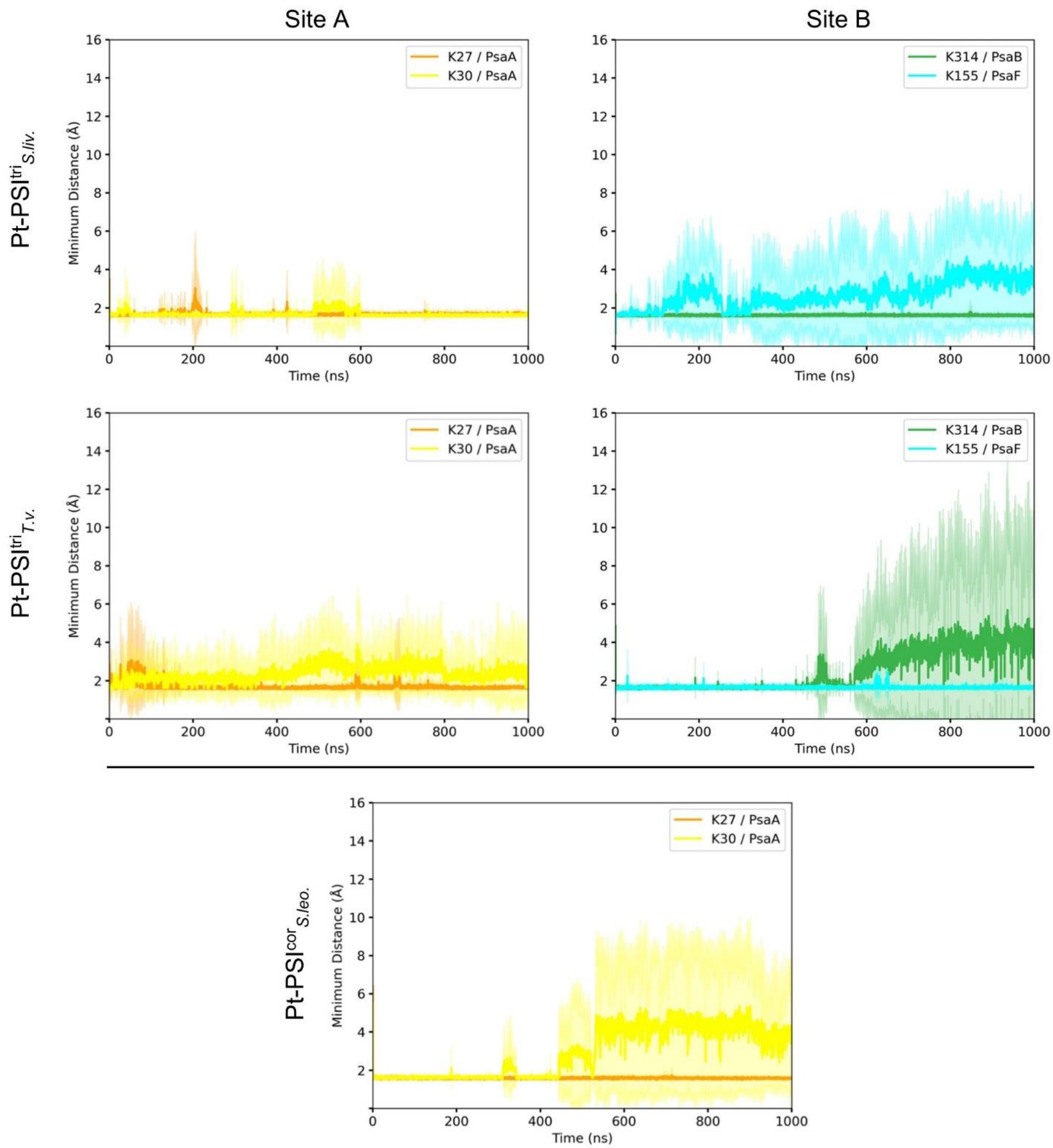


**Fig. S15.** Minimum distance between PtNP and key interacting residues of PSI. Time-resolved minimum distance (Å) between PtNP and two dominant interaction residues calculated over the 1 μs simulation. For the PtNPs in site A of the trimeric structures and the single site on the core structure, the distance over time is shown between the PtNP and PsaA-Lys27 and Lys30. For the PtNPs in site B of the trimeric structures, the distance over time is shown between the PtNP and PsaB-Lys314 and PsaF-Lys155.

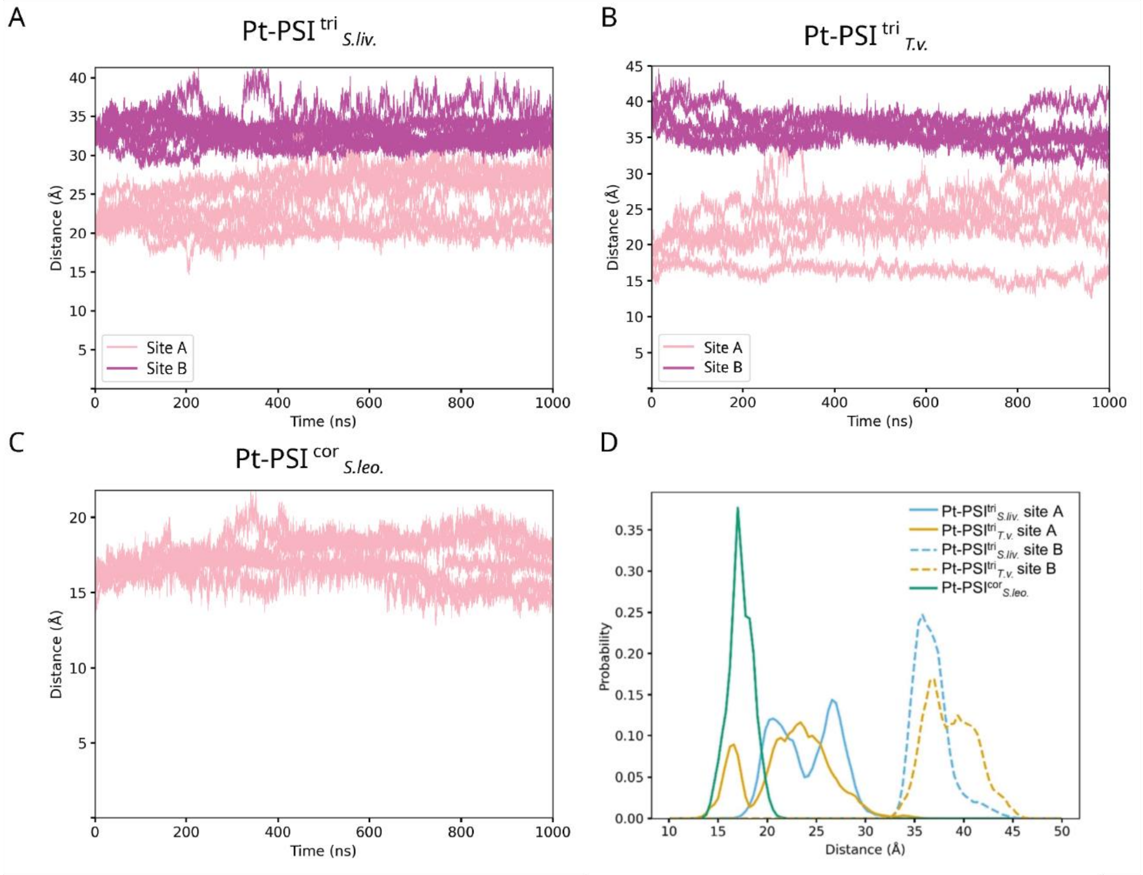


**Fig. S16.** Distance of PtNPs from the nearest [4Fe-4S] cluster in PSI-PtNP structures. The time-dependent minimum distance (*A*-*C*) is shown, with each line representing a single simulation trajectory. (*D*) Distance distributions for the nearest PtNP atom to the terminal [4Fe-4S] cluster in each PSI.

.

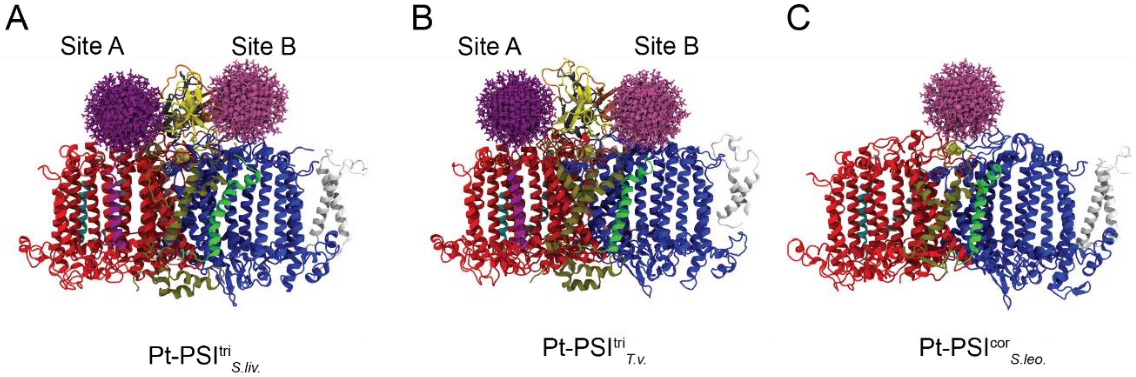


**Fig. S17.** Visual representations of the three PSI-PtNP biohybrid system coordinates used for MD. The site A and B nanoparticles are shown in pink and purple respectively, while the individual chains are shown in independently colored cartoon representations.

### Tables

**Table S1.** Sequence identities among PSI subunits from *S. leopoliensis*, *T. vestitus*, *S. lividus*, and *Synechocystis* 6803. Values are shown in units of percent and are colored red in a gradient from 0 (reddest) to 100 (white). PsaM is not included for *S. leopoliensis* because it is not found in the incomplete genome of *S. leopoliensis* despite it being clearly present in the cryo-EM map. Also note that of these organisms, PsaX is found only in *T. vestitus* and *S. lividus*. Sequence identities were calculated using Clustal Omega(6, 7).

| **PsaA** | *S. leopoliensis* | *T. vestitus* | *S. lividus* | *Synechocystis* 6803 |
|---|---|---|---|---|
| *S. leopoliensis* | | 86 | 86 | 85 |
| *T. vestitus* | 86 | | 98 | 88 |
| *S. lividus* | 86 | 98 | | 88 |
| *Synechocystis* 6803 | 85 | 88 | 88 | |

| **PsaB** | *S. leopoliensis* | *T. vestitus* | *S. lividus* | *Synechocystis* 6803 |
|---|---|---|---|---|
| *S. leopoliensis* | | 86 | 86 | 86 |
| *T. vestitus* | 86 | | 98 | 98 |
| *S. lividus* | 86 | 98 | | 88 |
| *Synechocystis* 6803 | 86 | 98 | 88 | |

| **PsaC** | *S. leopoliensis* | *T. vestitus* | *S. lividus* | *Synechocystis* 6803 |
|---|---|---|---|---|
| *S. leopoliensis* | | 95 | 95 | 98 |
| *T. vestitus* | 95 | | 100 | 95 |
| *S. lividus* | 95 | 100 | | 95 |
| *Synechocystis* 6803 | 98 | 95 | 95 | |

| **PsaD** | *S. leopoliensis* | *T. vestitus* | *S. lividus* | *Synechocystis* 6803 |
|---|---|---|---|---|
| *S. leopoliensis* | | 67 | 66 | 73 |
| *T. vestitus* | 67 | | 94 | 68 |
| *S. lividus* | 66 | 94 | | 71 |
| *Synechocystis* 6803 | 73 | 68 | 71 | |

| **PsaE** | *S. leopoliensis* | *T. vestitus* | *S. lividus* | *Synechocystis* 6803 |
|---|---|---|---|---|
| *S. leopoliensis* | | 70 | 74 | 74 |
| *T. vestitus* | 70 | | 87 | 68 |
| *S. lividus* | 74 | 87 | | 70 |
| *Synechocystis* 6803 | 74 | 68 | 70 | |

| **PsaF** | *S. leopoliensis* | *T. vestitus* | *S. lividus* | *Synechocystis* 6803 |
|---|---|---|---|---|
| *S. leopoliensis* | | 67 | 66 | 58 |
| *T. vestitus* | 67 | | 96 | 60 |
| *S. lividus* | 66 | 96 | | 59 |
| *Synechocystis* 6803 | 58 | 60 | 59 | |

| **PsaI** | *S. leopoliensis* | *T. vestitus* | *S. lividus* | *Synechocystis* 6803 |
|---|---|---|---|---|
| *S. leopoliensis* | | 58 | 55 | 55 |
| *T. vestitus* | 58 | | 95 | 68 |
| *S. lividus* | 55 | 95 | | 63 |
| *Synechocystis* 6803 | 55 | 68 | 63 | |

| **PsaJ** | *S. leopoliensis* | *T. vestitus* | *S. lividus* | *Synechocystis* 6803 |
|---|---|---|---|---|
| *S. leopoliensis* | | 73 | 67 | 68 |
| *T. vestitus* | 73 | | 93 | 68 |
| *S. lividus* | 67 | 93 | | 68 |
| *Synechocystis* 6803 | 68 | 68 | 68 | |

| **PsaK** | *S. leopoliensis* (PsaK1) | *S. leopoliensis* (PsaK2) | *T. vestitus* | *S. lividus* | *Synechocystis* 6803 |
|---|---|---|---|---|---|
| *S. leopoliensis* (PsaK1) | | 56 | 58 | 59 | 56 |
| *S. leopoliensis* (PsaK2) | 56 | | 49 | 52 | 44 |
| *T. vestitus* | 58 | 49 | | 93 | 48 |
| *S. lividus* | 59 | 52 | 93 | | 51 |
| *Synechocystis* 6803 | 56 | 44 | 48 | 51 | |

| **PsaL** | *S. leopoliensis* | *T. vestitus* | *S. lividus* | *Synechocystis* 6803 |
|---|---|---|---|---|
| *S. leopoliensis* | | 42 | 43 | 45 |
| *T. vestitus* | 42 | | 92 | 73 |
| *S. lividus* | 43 | 92 | | 71 |
| *Synechocystis* 6803 | 45 | 73 | 71 | 100 |

| **PsaM** | *S. leopoliensis* | *T. vestitus* | *S. lividus* | *Synechocystis* 6803 |
|---|---|---|---|---|
| *S. leopoliensis* | | 72 | 66 | 59 |
| *T. vestitus* | 72 | | 94 | 81 |
| *S. lividus* | 66 | 94 | | 77 |
| Synechocystis 6803 | 59 | 81 | 77 | |

| **PsaX** | *T. vestitus* | *S. lividus* |
|---|---|---|
| *T. vestitus* | | 97 |
| *S. lividus* | 97 | |

**Table S2.** Cryo-EM data collection, refinement, and validation statistics for trimeric *T. vestitus* PSI+PtNPs and core *S. leopoliensis* PSI+PtNP.

| | **Trimeric *T. vestitus* PSI+PtNPs** | **Core *S. leopoliensis* PSI+PtNP** |
|---|---|---|
| Magnification | x79,000 | x79,000 |
| Voltage (kV) | 200 | 200 |
| Electron exposure ($e^{-}Å^{-2}$) | 50.0 | 50.0 |
| Defocus range (µm) | −1.0 to −2.5 | −1.0 to −2.5 |
| Pixel size (Å) | 1.064 | 1.064 |
| Symmetry imposed | C3 | C1 |
| Initial particle images (no.) | 906,919 | 788,262 |
| Final particle images (no.) | 19,151 | 53,098 |
| Map resolution (Å) | 3.39 | 3.57 |
| FSC threshold | 0.143 | 0.143 |
| **Refinement** | | |
| Initial model used (PDB code) | 1JB0 | 1JB0 |
| Model resolution (Å) | 3.53 | 3.86 |
| FSC threshold | 0.5 | 0.5 |
| Map resolution range (Å) | 2.80-5.20 | 3.40-4.60 |
| Map-sharpening *B* factor ($Å^2$) | −89.3 | −109.8 |
| Model composition | | |
| Non-hydrogen atoms | 71,751 | 18,035 |
| Protein residues | 6,726 | 1,716 |
| Ligands | 381 | 105 |
| *B* factors ($Å^2$) | | |
| Protein | 66 | 187 |
| Ligands | 64 | 172 |
| R.m.s. deviations | | |
| Bond lengths (Å) | 0.012 | 0.007 |
| Bond angles (°) | 2.176 | 2.122 |
| **Validation** | | |
| MolProbity | 1.80 | 1.89 |
| Clashscore | 6.48 | 7.35 |
| Rotamer outliers (%) | 0.00 | 0.00 |
| Ramachandran plot | | |
| Favored (%) | 93.17 | 92.1 |
| Allowed (%) | 6.54 | 7.36 |
| Disallowed (%) | 0.29 | 0.55 |

**Table S3.** Interaction contributors to the PtNPs in Pt-PSI$^{tri}_{S.liv.}$. The top ten residues and lower-frequency contributors are shown in the table.

| Site A | | | Site B | |
|---|---|---|---|---|
| **Residue** | **Average Interaction frequency (%)** | | **Residue** | **Average Interaction frequency (%)** |
| PsaA-Lys30 | 213.5 | | PsaX-Thr9 | 232.7 |
| PsaA-Lys27 | 200.8 | | PsaB-Lys314 | 200.1 |
| PsaE-Arg39 | 163.9 | | PsaB-Lys408 | 153.2 |
| PsaE-Arg4 | 161.5 | | PsaF-Lys155 | 141.8 |
| PsaA-Arg36 | 131.7 | | PsaB-Arg414 | 122.2 |
| PsaA-Arg40 | 91.0 | | PsaX-Arg13 | 110.5 |
| PsaJ-Met1 | 64.6 | | PsaE-Arg9 | 107.2 |
| PsaE-Val2 | 54.5 | | PsaB-Lys308 | 74.0 |
| PsaJ-Lys4 | 36.8 | | PsaE-Lys7 | 33.3 |
| PsaF-Arg112 | 10.1 | | PsaF-Arg164 | 22.2 |
| **Top 10** | **1,128.5** | | **Top 10** | **1,197.1** |
| The rest | 35.2 | | The rest | 49.3 |

**Table S4.** Interaction contributors to the PtNPs in Pt-PSI$^{tri}_{T.v.}$. The top ten residues and lower-frequency contributors are shown in the table.

| Site A | | | Site B | |
|---|---|---|---|---|
| **Residue** | **Average Interaction frequency (%)** | | **Residue** | **Average Interaction frequency (%)** |
| PsaA-Lys27 | 121.0 | | PsaF-Lys155 | 117.9 |
| PsaA-Arg40 | 108.8 | | PsaB-Lys314 | 111.0 |
| PsaJ-Lys2 | 107.4 | | PsaF-Lys145 | 85.9 |
| PsaE-Arg4 | 92.2 | | PsaX-Arg16 | 49.5 |
| PsaA-Lys30 | 90.9 | | PsaX-Pro11 | 35.7 |
| PsaA-Arg36 | 74.2 | | PsaE-Lys9 | 28.6 |
| PsaJ-Met1 | 56.5 | | PsaE-Lys7 | 25.0 |
| PsaF-Lys120 | 54.7 | | PsaB-Lys408 | 13.3 |
| PsaE-Arg40 | 51.9 | | PsaF-Asn157 | 12.2 |
| PsaC-Lys35 | 28.5 | | PsaE-Arg4 | 9.2 |
| **Top 10** | **786.1** | | **Top 10** | **488.2** |
| The rest | 23.3 | | The rest | 42.2 |

**Table S5.** Interaction contributors to the PtNPs in Pt-$PSI^{cor}_{S.leo.}$. The top ten residues and lower-frequency contributors are shown in the table.

| Residue | Average Interaction frequency (%) |
|---|---|
| PsaA-Lys27 | 290.8 |
| PsaB-Arg564 | 239.5 |
| PsaA-Lys40 | 236.6 |
| PsaB-Lys545 | 216.4 |
| PsaA-Lys30 | 174.8 |
| PsaA-Lys43 | 160.1 |
| PsaF-Lys110 | 131.8 |
| PsaA-Arg583 | 131.3 |
| PsaA-Arg728 | 124.3 |
| PsaA-Lys720 | 96.8 |
| **Top 10** | **1802.4** |
| The rest | 376.6 |

**SI References**


1. O. Carugo, How root-mean-square distance (r.m.s.d.) values depend on the resolution of protein structures that are compared. *J. Appl. Crystallogr.* **36**, 125–128 (2003).
2. A. Einstein, Über die von der molekularkinetischen Theorie der Wärme geforderte Bewegung von in ruhenden Flüssigkeiten suspendierten Teilchen. *Ann. Phys.* **322**, 549–560 (1905).
3. F. B. Sheinerman, C. L. Brooks, Molecular picture of folding of a small α/β protein. *Proc. Nat'l Acad. Sci.* **95**, 1562–1567 (1998).
4. M. Ceriotti, J. Cuny, M. Parrinello, D. E. Manolopoulos, Nuclear quantum effects and hydrogen bond fluctuations in water. *Proc. Nat'l Acad. Sci.* **110**, 15591–15596 (2013).
5. L. B. Skinner, C. J. Benmore, J. C. Neuefeind, J. B. Parise, The structure of water around the compressibility minimum. *J. Chem. Phys.* **141**, 214507 (2014).
6. F. Sievers, D. G. Higgins, Clustal Omega for making accurate alignments of many protein sequences. *Protein Sci.* **27**, 135–145 (2018).
7. F. Sievers, *et al.*, Fast, scalable generation of high-quality protein multiple sequence alignments using Clustal Omega. *Mol. Syst. Biol.* **7**, 539 (2011).
